# Altering CLC stoichiometry by reducing non-polar side-chains at the dimerization interface

**DOI:** 10.1101/2020.10.29.361279

**Authors:** Kacey Mersch, Tugba N. Ozturk, Kunwoong Park, Hyun-Ho Lim, Janice L. Robertson

## Abstract

CLC-ec1 is a Cl^-^/H^+^ antiporter that forms stable homodimers in lipid bilayers, with a free energy of −10.9 kcal/mole relative to the 1 subunit/lipid standard state in 2:1 POPE/POPG lipid bilayers. The dimerization interface is formed by four transmembrane helices: H, I, P and Q, that are lined by non-polar side-chains that come in close contact, yet it is unclear as to whether their interactions drive dimerization. To investigate whether non-polar side-chains are required for dimer assembly, we designed a series of constructs where side-chain packing in the dimer state is significantly reduced by making 4-5 alanine substitutions along each helix (H-ala, I-ala, P-ala, Q-ala). All constructs are functional and three purify as stable dimers in detergent micelles despite the removal of significant side-chain interactions. On the other hand, H-ala shows the unique behavior of purifying as a mixture of monomers and dimers, followed by a rapid and complete conversion to monomers. In lipid bilayers, all four constructs are monomeric as examined by single-molecule photobleaching analysis. Further study of the H-helix shows that the single mutation L194A is sufficient to yield monomeric CLC-ec1 in detergent micelles and lipid bilayers. X-ray crystal structures of L194A reveal the protein re-assembles to form dimers, with a structure that is identical to wild-type. Altogether, these results demonstrate that non-polar membrane embedded side-chains play an important role in defining dimer stability, but the stoichiometry is highly contextual to the solvent environment. Furthermore, we discovered that L194 is a molecular hot-spot for defining dimerization of CLC-ec1.

## INTRODUCTION

The cell membrane encapsulates all living things with a structurally defined layer of oil - the lipid bilayer. This hydrocarbon core provides an electrostatic barrier to the passive permeation of ions and charged molecules. With that, biology has evolved a special class of membrane proteins, that stably reside within the lipid bilayer and control the precise transport of chemicals and information across the membrane barrier. Membrane proteins are built with the same amino acids that are found within water soluble proteins. However, they must be lined by hydrophobic amino acids at their lipid exposed surfaces to enable partitioning into the lipid bilayer while not disrupting the integrity of the membrane. In the past, it was proposed that membrane proteins have evolved as inversions of soluble proteins, favoring polar and hydrophilic interactions within the core, while exposing the non-polar surfaces to the surrounding lipids (Engelman and Zaccai 1980).

While it is true that membrane proteins display hydrophobic residues on the lipid-facing surfaces, and that they often contain hydrophilic permeation or transport pores within, it is not generally true that membrane proteins interact via polar groups embedded within the membrane. In fact, many membrane protein complexes assemble via non-polar surfaces, between transmembrane helix contacts in multi-helix folds, or along larger surfaces that are involved in oligomerization (Oberai et al. 2009). In these cases, the hydrophobicity of the interaction interfaces appears to be similar to that of lipid-facing interfaces. This raises the question of what drives the assembly of membrane proteins via these greasy interfaces, when they appear to be suitably solvated by the similarly greasy lipids in the membrane?

One possibility is that non-polar side-chains at these interfaces allow specific van der Waals (VDW) interactions to form, which exclusively stabilize the assembled state. While some degree of VDW interactions is also expected between the exposed interfaces and the lipids in the dissociated state, it may be that the protein-protein interactions in the associated complex are optimized leading to a net stabilizing driving force for assembly. Indeed, VDW interactions by side-chains have been shown to play an important role in the dimerization stability of Glycophorin-A (Doura et al. 2004), however in this system, it has also been shown that other backbone contributions, such as hydrogen bonding also contribute to the overall stability (MacKenzie, Prestegard, and Engelman 1997; Senes, Engel, and DeGrado 2004). Thus, it remains an open question whether side-chains at non-polar surfaces are indeed responsible for driving the assembled state over the dissociated lipid solvated states.

Recently, we have developed a new type of model system that has the potential to expand our ability to investigate how interfacial non-polar side-chains affect membrane protein assembly. CLC-ec1 is a homodimeric Cl^-^/H^+^ antiporter that is native to *E. coli* inner membranes (**Figure 1A**). It is a large protein, with 18 transmembrane helices per subunit (A-R) that associate to form a stable dimer, *ΔG*° = −10.9 kcal/mole, 1 subunit/lipid standard state, in 2:1 POPE/POPG lipid bilayers measured using single-molecule photobleaching analysis (Chadda et al. 2018; Chadda et al. 2016; Chadda and Robertson 2016). The dimerization interface is large, comprised of four transmembrane helices (H, I, P & Q) with a surface area of ~1200 Å^2^ lined by ≈ 20 non-polar residues **(Figure 1B)**. These surfaces exhibit high shape complementarity placing all of these side-chains within VDW contact distance (Robertson, Kolmakova-Partensky, and Miller 2010).

**Figure 1.**
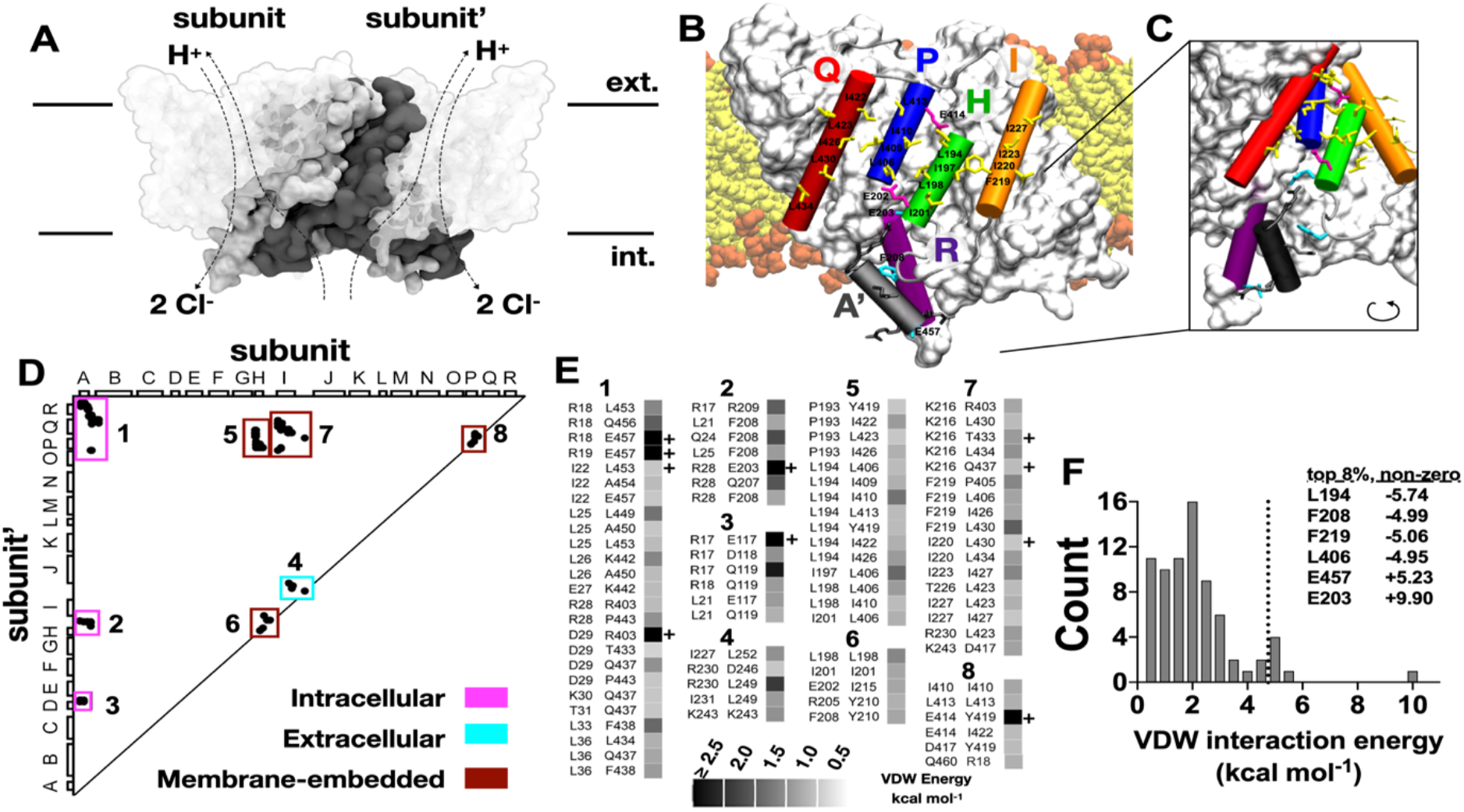
Computational analysis of side-chain interactions at the dimerization interface of CLC-ec1. (A) Homodimeric CLC-ec1 Cl^-^/H^+^ antiporter (PDB ID: 1OTS). The protein surface comprising the dimerization interface, all residues within 5 Å of the adjacent subunit, is highlighted in opaque surface representation. (B) The dimerization interface is comprised by four membrane embedded helices shown in cartoon representation: Q - red, P - blue, H - green and I - orange, following the 18 transmembrane A-R helix nomenclature (Dutzler et al., 2002). Side-chains involved in dimerization are highlighted in licorice representation. (C) Close-up and rotated view of the cytosolic contact surface between helix R - purple and helix A’ - black, on the adjacent subunit’. (D,E) Pair-wise side-chain VDW interaction energies E_VDW_ > |0.5 kcal/mole| based on the energy minimized 1OTS structure. Pairs highlighted as “+” represent VDW clashes that persist after minimization. (F) Histogram of E_VDW_ for the interaction between a residue on one subunit with all residues on the opposing subunit’. The dotted line marks the 92^nd^ percentile of all interactions > |0.25 kcal/mole|, and the top contributors are listed in the table.

To investigate whether side-chains are important for defining dimerization stability, we carried out an extreme mutagenesis study where we strip each helix of all interfacial side-chains, making 4-5 simultaneous substitutions to alanine (8-10 in the dimer state). These “helix-ala” constructs are predicted to form cavities within the dimerization interface, resulting in a significant loss of side-chain interactions. By examining monomer-dimer populations in detergent and lipid bilayers, carrying out functional transport assays and examining structural changes by x-ray crystallography, we are able to determine that these side-chains contribute to the overall dimerization stability in lipid bilayers, but that they are not required for dimerization in detergent micelles. Furthermore, we identify that residue L194 participates in a molecular hot-spot for CLC-ec1 dimerization. Altogether, these results demonstrate that interfacial side-chains impact dimer stability in lipid bilayers, and that membrane protein stoichiometry is a highly contextual observable that depends strongly on the solvent environment.

## RESULTS

### Computational analysis of protein interactions at the dimerization interface of CLC-ec1

The CLC-ec1 dimer exhibits many side-chain contacts between the two subunits, all of which may contribute stabilizing interactions in the dimer state (**Figure 1A**). Visually, these contacts are found along four membrane embedded helices H, I, P & Q, between the domain-swapped helix A, the C-terminal helices Q & R, and the extracellular loop between helices I & J (**Figure 1B, C**). To quantify these contributions, we calculated the pair-wise VDW interaction energy between side-chains of opposing subunits, designated subunit & subunit’, in the energy minimized PDB ID: 1OTS dimer structure (Dutzler, Campbell, and MacKinnon 2003) (**Figure 1D, E, Supplementary Table 1**). Absolute energy values > 0.5 kcal/mole, were categorized further into intracellular, extracellular and membrane embedded regions. Intracellular interactions involve the N-terminal helix A with (1) helices Q’/R’, (2) the H’-I’ loop and (3) the D’-E’ loop. On the extracellular side, there are interactions between (4) the I-J and I’-J’ loops. The remaining contacts are all membrane-embedded and occur between (5) helix H and helices P’/Q’, (6) helix H and helix H’, (7) helix I and helices P’/Q’, and (8) helix P with helices P’/Q’.

The total VDW interactions that each residue makes with the opposing subunit was analyzed and the non-zero values plotted as a histogram (**Figure 1F, Supplementary Table 2**). This shows that the majority of interactions are weak, with the majority contributing ≈ 2 kcal/mole. However, there is a rightward tail indicating residues with increased packing relative to the rest of the interface. The top 8% of contributors include three cytosolic residues E203 and F208 that occur at the end, or just after helix H, and E457 on helix R. All three of these side-chains exhibit anomalous packing interactions with the N-terminal helix A’ of the opposing subunit that undergoes a domain-swapping configuration **(Figure 1C)**. Note, two of these residues, E203 and E457, exhibit positive VDW interaction energies corresponding to steric clashes. Despite the significant number of contacts involving helix A’, previous studies showed that the crystal structure of the N-terminal helix A truncated construct yields the dimeric form of the protein, indicating that helix A is not required for dimerization (Lim, Shane, and Miller 2012). The other three significant contributors to VDW interactions are considered membrane embedded, and include L194, F219 and L406. Since phenylalanine is the largest non-polar side-chain, it is not surprising that F219 stands out as a stronger contributor. However, the other two sites are occupied by leucine, and so their appearance indicates that the packing at these positions is significantly increased relative to the other regions on the interface. It is important to note, that L194 and L406 represent homologous sites in the inverted topology fold of this protein (Dutzler et al. 2002), both lining the symmetry interface that occurs between helices H & P. With the exception of F219, the remaining five strongest cytosolic and membrane embedded VDW contributors are all generally located along this symmetry interface, indicating an increase in side-chain packing related to the inverted topology fold.

### Dimerization of helix-ala CLC-ec1 constructs in detergent micelles

From here, we focused on investigating whether membrane embedded VDW interactions in the dimer state are required for CLC-ec1 dimerization. We selected the major VDW contributors across each helix and calculated their cumulative side-chain interactions with the opposing subunit (**Figure 2A**). Removal of these side-chains along each helix is predicted to yield a 10-14 kcal/mol loss of interactions in the dimer state per subunit. Following this, we designed four CLC-ec1 constructs in which 4-5 residues were mutated to alanine along each of the dimerization helices, i.e. “helix-ala”. We refer to these specifically as H-ala: L194A/I197A/L198A/I201A, I-ala: F219A/I220A/I223A/I227A, P-ala: L406A/I409A/I410A/L413A and Q-ala: I422A/L423A/I426A/L430A/L434A (**Figure 2B**). Calculation of the cavity volume indicates that large 350-450 Å^3^ pockets are formed at the dimerization interface in the helix-ala constructs provided the backbone structure in the dimer complex does not change (**Figure 2C**). Based on this, we hypothesize that these constructs will yield a significant destabilization of the dimer complex and yield monomeric protein.

**Figure 2.**
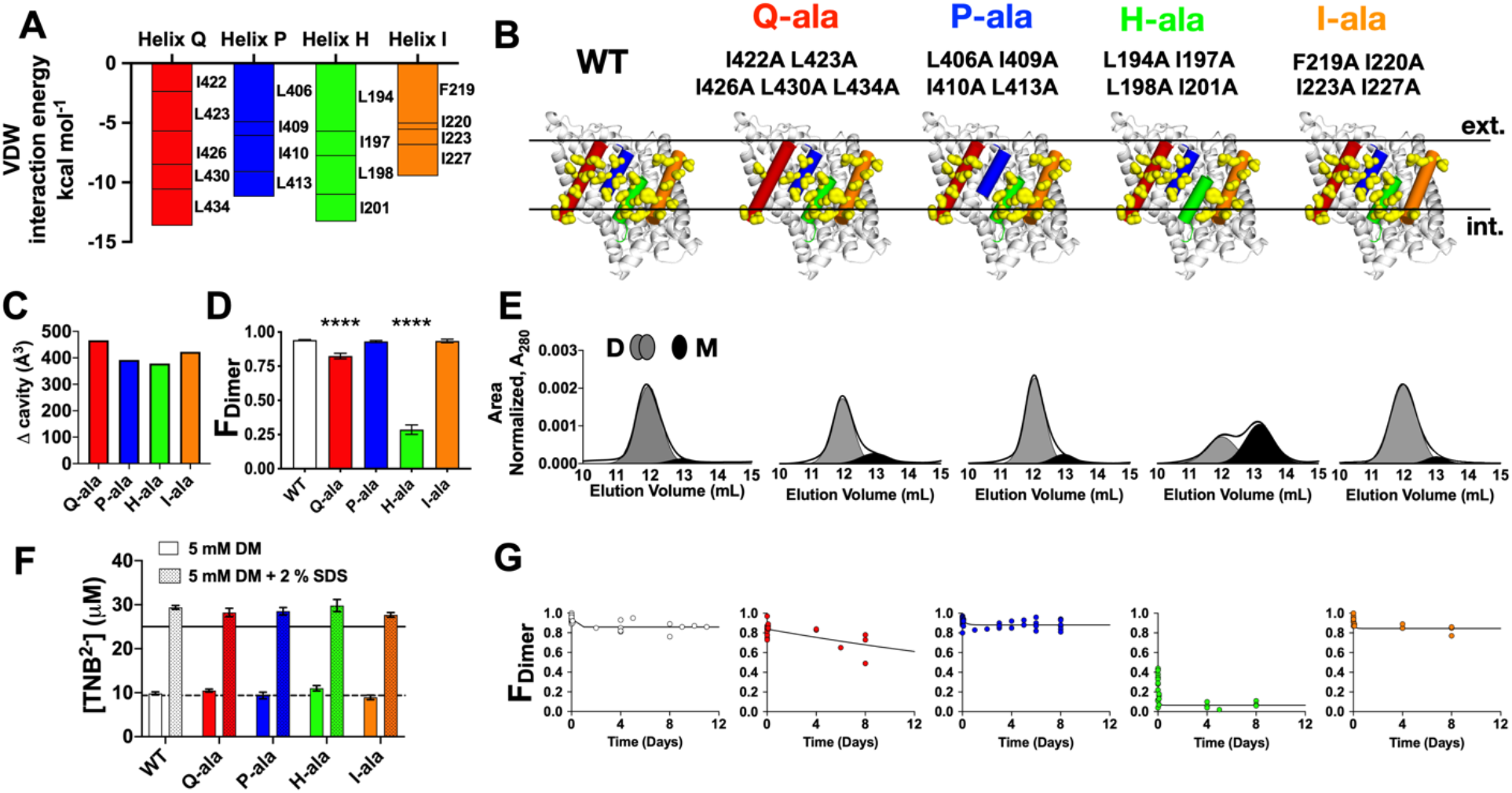
The effect of removing membrane embedded side chain contacts on CLC-ec1 dimerization in detergent micelles. (A) Sum of VDW interaction energies for the membrane embedded portion along each dimerization interface helix. Contributions from each individual residue are listed. (B) The dimerization interface of CLC-ec1 is shown (WT), with membrane embedded side chain contacts highlighted (yellow surface). Four constructs were designed to reduce side-chain interactions in the dimer state by simultaneous mutation of 4-5 residues to alanine along each interfacial helix: ‘Q-ala’ – I422A/L423A/I426A/L430A/L434A, ‘P-ala’ – L406A/I409A/I410A/L413A, ‘H-ala’ – L194A/I197A/L198A/I201A, and ‘I-ala’ – F219A/I220A/I223A/I227A. (C) Difference in cavity volumes, with respect to the cavity volume of the wild-type structure, was calculated for the H-ala, I-ala, P-ala and Q-ala structures. The volumes correspond to the cavity formed in the dimer state. (D) Fraction of dimeric protein, *F_Dimer_*, from the initial purification from *E. coli*. (E) SEC profiles of the purified CLC-ec1 constructs in 5 mM DM micelles. (F) Cysteine accessibility reaction of C85A/H234C CLC-ec1 with Ellman’s reagent (DNTB) as a measure of protein folding in detergent. Data represent mean ± sem; sample n_wT_ = 11, n_Q-ala_ = 4, n_P-ala_ = 9, n_H-ala_ = 3, n_I-ala_ = 4. The multiple alanine variants show no significant difference compared with WT (solid - 5 mM DM and patterned - 5 mM DM + 2% SDS). (G) Fraction of dimeric protein, *F_Dimer_*, calculated from SEC for the initial purification (t = 0) and subsequent re-runs of the protein following incubation at 10 μM and at 4°C, χ = 3 x 10^−3^ subunits/detergent. Data reflect independent purifications, n_WT_ = 6, n_Q-ala_ = 3, n_P-ala_ = 12, n_H-ala_ = 4, n_I-ala_ = 4 was, and is fit with an exponential decay function.

To test this hypothesis, wild-type (WT) CLC-ec1 and the helix-ala constructs were expressed in *E. coli* and purified in buffer containing 5 mM n-Decyl-β-D-Maltopyranoside (DM) detergent (Chadda et al. 2016). Size exclusion chromatography (SEC) analysis of the protein in detergent micelles shows that I-ala, P-ala and Q-ala are dimeric upon purification, whereas H-ala shows a mixture of monomers and dimers. (**Figure 2D, E**). To examine whether the monomeric shift for H-ala reports on a change in subunit fold, we carried out a cysteine accessibility experiment using Ellman’s reagent **(Figure 2F)**. In this assay, WT CLC-ec1, on the background of C85A/H243C, has one water-accessible cysteine that is used for rapid conjugation by fluorophore-maleimides, and two native buried cysteines, C302 and C347, that are only reactive when the protein is denatured by sodium dodecyl sulfate (SDS) (Chadda et al. 2016). All of the helix-ala constructs, including H-ala, show a single accessible cysteine in DM micelles, and three in SDS, indicating that they all share a similar fold to WT in DM micelles. Next, we investigated whether the monomer-dimer distribution changes with time, examining the protein by SEC over 12 days, incubating at 4 °C and at a constant protein to detergent molar ratio, 3 x 10^−3^ subunits/detergent (**Figure 2G**). P-ala and I-ala show no shift in dimerization compared to WT, while Q-ala shows a possible slow decay (t1/2 = 26 days). On the other hand, H-ala rapidly converts to an all-monomeric population by the first re-run on the column (t = 54 ± 4 minutes, mean ± SEM). Therefore, on a reasonable laboratory timescale (time > 1 week), the helix-ala constructs remain dimeric in DM micelles, with the exception of H-ala which is uniquely monomeric.

### The stoichiometry of CLC-ec1 helix-ala constructs in 2:1 POPE/POPG lipid bilayers

Next, we investigated the behavior of the CLC-ec1 helix-ala constructs in membranes. To assess the functionally relevant folded state, we measured the chloride transport activity of Cy5 labelled helix-ala CLC-ec1 in 2:1 POPE/POPG lipid bilayers, where Cy5 labeling is required for later quantification of the stoichiometry in membranes. Chloride efflux was measured for proteoliposomes reconstituted at χ_reconst_. = 1.5 x 10^−5^ subunit/lipid, corresponding to 1 μg/mg protein/lipid (**Figure 3A**). All of the constructs showed chloride transport rates, *k_CLC_* and fractional volume of inactive vesicles, *F_0,Cl_*, that were significantly active compared to empty vesicles. However, H-ala-Cy5 and I-ala-Cy5 showed a small reduction in transport rate compared to WT-Cy5 (**Figure 3B**), and there was an increase in inactive population for H-ala-Cy5 and P-ala-Cy5 (**Figure 3C**). Note that the Cy5 labeling yield was similar for all constructs studied (**Figure 3D**). Therefore, while it can be concluded that all of the constructs are functional, the helix-ala constructs lead to slightly lower activity, with the most significant impact on H-ala-Cy5.

**Figure 3.**
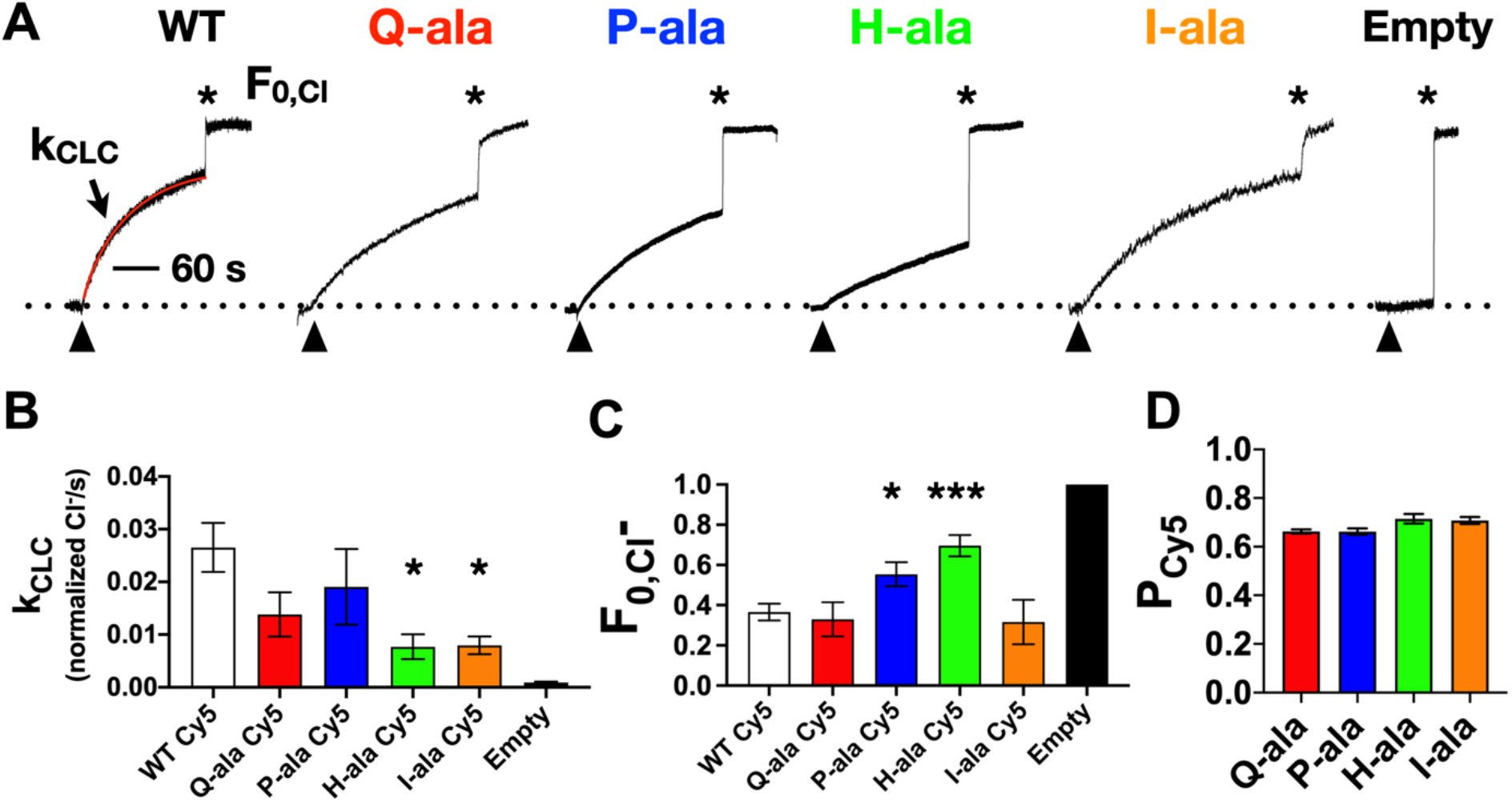
Chloride transport activity for helix-ALA CLC-ec1 constructs. (A) Normalized chloride efflux traces of 1 μg/mg (*χ*_reconstituted_ = 1.5 x 10^−5^ subunits/lipid) CLC-ec1-Cy5 in 2:1 POPE/POPG liposomes, extruded through a 0.4 μm nucleopore membrane after 7 freeze/thaw cycles. Triangles represent the initiation of chloride efflux with the addition of valinomycin (0.5 nM) and FCCP (1 nM), asterisks indicate the addition of β-OG to a final concentration of 30 mM, allowing for the measurement of the fractional volume of inactive vesicles, *F_0,Cl-_*. (B) Chloride efflux rate, *k_CLC_*, from the fit of the transport. (C) Fractional volume of inactive liposomes, *F_0,Cl-_*. Data represented as mean ± sem, n_WT_ = 12, n_Q-ala_ = 4, n_P-ala_ = 7, n_H-ala_ = 4, n_I-ala_ = 4. Significance was measured compared to WT transport rate or the rate of empty liposome leakage using a student t-test, *, *p* ≤ 0.05, and ***, *p* ≤ 0.001. Each protein measured in B has chloride transport rates significantly greater than Empty vesicles. (D) Cy5 labeling yield, *P_Cy5_*, per subunit for each construct. Data represented as mean ± sem, n_Q-ala_ = 4, n_P-ala_ = 7, n_H-ala_ = 4, n_I-ala_ = 4. The only differences found were that of P-ala and H-ala groups.

Next, we examined the dimerization reaction for the helix-ala constructs in 2:1 POPE/POPG lipid bilayers. For this, Cy5 labelled CLC-ec1 was reconstituted as a function of the protein to lipid mole fraction, χ_reconst_. = 2 x 10^−8^ to 5 x 10^−6^ subunits/lipid, the samples were imaged by total internal reflection fluorescence (TIRF) microscopy (**Figure 4A**) in order to carry out single-molecule photobleaching analysis (**Figure 4B**) and quantify the subunit-capture statistics. This method was previously used to quantify the Poisson-like distribution of protein reconstitution into liposomes from large equilibrium membranes, allowing for the measurement of the reversible WT CLC-ec1-Cy5 dimerization reaction with *ΔG*° = −10.9 kcal/mol relative to the 1 subunit/lipid standard state in 2:1 POPE/POPG lipid bilayers (Chadda et al. 2016; Chadda et al. 2018). Comparing the photobleaching probability distributions of the helix-ala constructs as a function of mole fraction (**Figure 4C-E**) shows that they behave similarly to I201W/I422W, ‘WW’, the monomeric control (Chadda et al. 2018). Calculating the fraction of dimer, *F_Dimer_*, by least-squares fitting of the helix-ala distribution to a linear combination of the monomer and dimer control distributions, shows that there is minimal dimerization within the protein density range studied (**Figure 4F**). While the reaction data is limited, fits of *F_Dimer_* to the dimerization isotherm provides a lower limit on the change of stability, with the helix-ala constructs destabilized by *ΔΔG* > +3.5 kcal/mol compared to WT. The actual destabilization is possibly larger, but the higher densities required to measure the reaction fall outside of the dynamic range of the single-molecule subunit capture approach. Still, these results demonstrate that the helix-ala constructs are monomeric in the 2:1 POPE/POPG lipid bilayers over the density range studied, indicating that the complexes are substantially destabilized while the monomers remain functionally competent for chloride transport.

**Figure 4.**
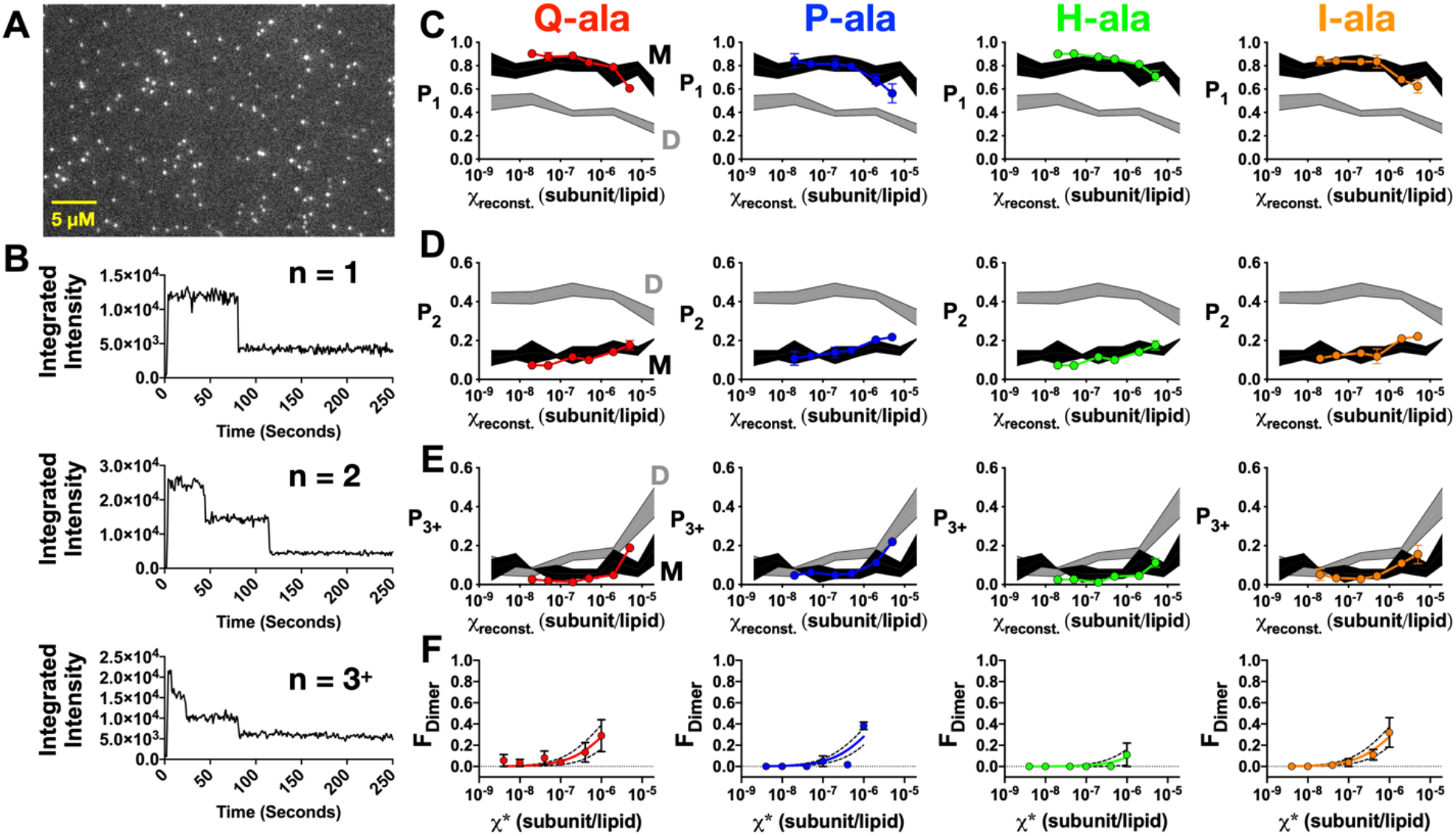
Single-molecule photobleaching analysis of the oligomeric distribution of helix-ala CLC-ec1 constructs in 2:1 POPE/POPG lipid bilayers. (A) Total internal reflection fluorescence microscopy image of Q-ala CLC-ec1-Cy5 liposomes at a χ_reconst_. = 5 x 10^−7^ (subunits/lipid). (B) Representative photobleaching traces single, double and three or more photobleaching steps corresponding to the *P*_1_, *P*_2_, *P*_3+_ populations in the photobleaching probability distribution. (C) *P*_1_, (D) *P*_2_, and (E) *P*_3+_ photobleaching probabilities as a function of the reconstituted mole fraction, χ_reconst_ (subunits/lipid). Data represent mean ± sem, n = 3. CLC-ec1 monomer (M – black, I201W/I422W) and dimer (D – grey, R230C/L294C) control data (Chadda et al., 2018) are shown for reference (mean ± standard error bands). (F) *F_Dimer_* as a function of the reactive mole fraction, *χ\** (subunits/lipid). Solid lines represent lower-limit fits to the dimerization isotherm, while dotted line indicate 95% CI bands. Data shown as mean ± sem, n = 3. Experiments were carried out in dialysis buffer (DB).

### Helix H contains a hot-spot for dimerization

While all of the helix-ala constructs were monomeric in lipid bilayers, H-ala was the only one to show a shift to monomers while in detergent micelles. To investigate this further, we constructed single (L194A, I197A, L198A, I201A), double (I197A/L198A), and triple (L194A/I197A/L198A) alanine substitutions on helix H (**Figure 5A**) and quantified the proportion of dimers in detergent upon purification by SEC (**Figure 5B**). All of these constructs were dimeric upon initial purification, with the exception of L194A and L194A/I197A/L198A. Since the common factor between these two constructs was L194A, we focused on this mutant for the remaining studies. L194A incubated at 4 °C or at room temperature (RT) at a mole fraction of 3 x 10^−3^ subunits/detergent, shows a change in the monomer-dimer population as examined by SEC (**Figure 5G**). Over several days, L194A decays into a stationary mixture of monomers and dimers, with *F_Dimer,4°C_* = 0.61 (t_1/2_ = 1.1 days) and *F_Dimer,RT_* = 0.53 (t1/2 = 0.4 days), compared to *F_Dimer_* = 0.85 for WT. Despite the increase in monomer over time, L194A demonstrates the same cysteine accessibility as WT, suggesting that the subunit fold is intact (**Figure 5C**). To examine whether L194A yields monomers due to a change in subunit structure, we determined its x-ray crystallography. C85A/L194A and C85A/H234C/L194A CLC-ec1 were purified and bound to the F_ab_ antibody chaperone and the structure of the complex was solved by x-ray diffraction at 3.0 and 2.9 Å resolution respectively. Both structures show that L194A forms dimers identical to the WT (**Figure 5E**) with no observable change at the dimerization interface (**Figure 5F**).

**Figure 5.**
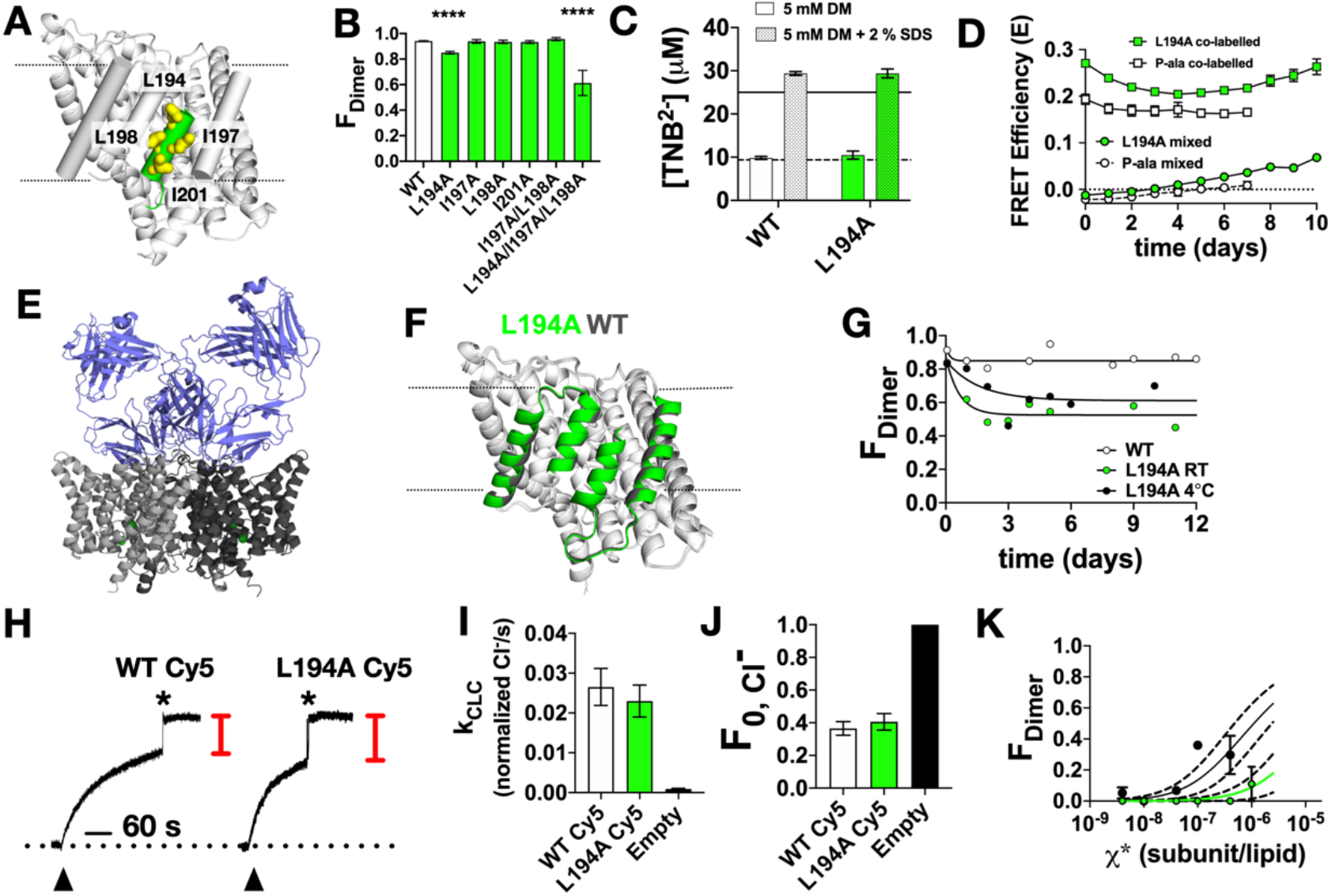
L194A disrupts dimerization in detergent micelles and lipid bilayers. (A) Close-up image of helix-H and residues involved in dimerization (yellow surface). (B) *F_Dimer_* for CLC-ec1 helix H single, double and triple alanine substitutions upon initial purification in 5 mM DM analyzed by SEC. Data represented as mean ± sem, for n_WT_ = 37, n_L194A_ = 49, n_I197A_ = 6, n_L198A_ = 6, n_I201A_ = 5, n_I197A/L198A_ = 3, n_I194A/L198A/I201A_ = 3, **** = *p* ≤ 0.001. (C) Cysteine accessibility of CLC-ec1 and L194A in 5 mM DM (dashed bar) and 5 mM DM + 2% SDS (solid bar). Data represent mean ± sem, n_WT_ = 11, n_L194A_ = 9. (D) FRET analysis of the possibility of L194A-Cy3/DM, L194A-Cy5/DM, P-ala-Cy3/DM, and P-ala-Cy5/DM subunit-exchange in 5 mM DM, showing no change of the FRET signals as a function of protein incubation at room temperature. FRET represents donor quenched bulk FRET, *E* = 1-*F_AD_*/*F_D_*, ‘mixed’ refers to combining protein after independently labeling with either Cy3 or Cy5, whereas ‘co-labelled’ refers to the positive control of either L194A or P-ala that was labelled with Cy3 and Cy5 simultaneously. Experiments were carried out in 150 mM Sodium Chloride (RPI), 20 mM MOPS, 5 mM DM buffer, pH of 7.0. (E) X-ray crystal structure of the CLC-ec1 L194A homodimer (PDB ID: 7CVS, 7CVT). The CLC-ec1 variants subunits are gray and white while the Fab is colored in slate. (F) An alignment between backbone only atoms of a single subunit of PDB ID: 1OTS (gray) and the L194A (green) structure. The parts of the structure outside of the Q, P, H, and I helices are colored white. The calculated RMSD was 0.5 Å. (G). SEC measured *F_Dimer_* for WT (white circles) at 10 μM and 5 mM DM (*χ* = 3 x 10^−3^ subunits/detergent molecule) and L194A incubated at 4°C (black circles) and RT (green circles) mean ± sem, for n_WT_ = 6, n_L194A,4°C_ = 7, n_L194A,RT_ = 5. Data fit to exponential decay: *k_WT_* = 5.85 days^−1^, plateau=0.85, *k_L194A,4°C_* = 0.63 days^−1^, plateau = 0.61, *k_L194A,RT_* = 1.56 days^−1^, plateau = 0.53. (H) Normalized chloride efflux traces of 1 μg/mg (*χ_reconst._* = 1.5 x 10^−5^ subunit/lipid) CLC-ec1-Cy5 in 2:1 POPE/POPG liposomes, extruded through a 0.4 μm nucleopore membrane after 7 freeze/thaw cycles. Triangles represent the initiation of chloride efflux with the addition of valinomycin (0.5 nM) and FCCP (1 nM), asterisks indicate the addition of β-OG to a final concentration of 30 mM, allowing for the measurement of the fractional volume of inactive vesicles, *F_0,Cl-_*. (I) Fraction of inactive vesicle volume, *F_0,Cl_-* and (J) chloride efflux rate, *k_CLC_*, for CLC-ec1 H-helix single-alanine substitutions in 2:1 POPE/POPG 400 nm extruded liposomes. Data represented as mean ± sem, for n_WT_ = 12, n_L194A_ = 5. (K) *F_Dimer_* of L194A in 2:1 POPE/POPG membranes measured by the single-molecule photobleaching subunit capture method. Line represents the best-fit ± errors to a dimerization isotherm, indicating a lower limit of the *K_D,L194A_* = 1.1 x 10^−6^ subunit/lipid. Data represent mean ± sem, n = 5. H-ala data was added here as a “monomeric reference” for comparison with L194A which is undergoing a dimerization reaction.

The relaxation of the protein into a stationary population of monomers and dimers **(Figure 5G)** and the reassembly of the dimer during crystallization (**Figure 5E, F**) presents a possibility that L194A may participate in a dynamic dimer equilibrium in detergent micelles. This behavior has been observed previously for the F^-^/H^+^ antiporter CLC^F^-eca, a homologue of CLC-ec1, in DM micelles (Last and Miller 2015). To test whether this is the case, we first examined whether the addition of DM micelles shifts the system towards monomers, studying L194A and P-ala in parallel (**Supplemental Figure 3A**). For either construct, addition of detergent did not yield a clear shift; however, it is possible that the extent of dilution was insufficient based on the reaction equilibrium or that the kinetics for equilibration of the system was beyond our observation time. To further investigate this, we applied an alternate approach to examine whether these protein species showed evidence for dynamic subunit exchange in the DM micelles by measuring the formation of a heterodimeric CLC-ec1 species that is capable of participating in Förster Resonance Energy Transfer (FRET). P-ala or L194A were labelled individually with either Cy3 or Cy5, and then mixed together and examined alongside a co-labelled Cy3/Cy5 sample, representing the end-point positive control. The FRET efficiency was measured over the course of 6 days while incubating at room temperature (≈ 22 °C) (**Figure 5D**). Over this time, we observed no increase in FRET for the mixed samples, and a stable FRET signal for the co-labelled controls. After one week, we observed increases in FRET in the L194A samples that may indicate non-specific aggregation, as we found similar increases in our control samples when incubated at 37 °C (**Supplemental Figure 3B**). However, the lack of increase in FRET for the mixed samples over the relaxation timescale exhibited in the SEC analysis (**Figure 5G**) suggests that the stationary mixture of L194A does not reflect a steady-state equilibrium, but is instead an irreversible decay in the current SEC conditions. Thus, the ability for L194A, and H-ala to convert to monomers indicates an ability for these constructs to destabilize and rupture the micelle.

Finally, we examined the behavior of the L194A CLC-ec1 in lipid bilayers. Measurements of the functional activity of L194A (**Figure 5H**) shows no change in the fraction of inactive vesicles or the chloride transport rate compared to WT (**Figure 5I, J**). We then examined the dimerization reaction of L194A in 2:1 POPE/POPG lipid bilayers using the single-molecule photobleaching subunit capture approach (**Supplementary Figure 3C-F**). We found that L194A was significantly destabilized in membranes (**Figure 5K**), and could observe the early phase of a dimerization reaction. Fitting the limited association data to a dimerization isotherm yields a lower limit estimation of the destabilization of *ΔΔG_L194A_* > +2.9 kcal/mol compared to WT. Thus, the single mutation of L194A alone yields monomeric CLC-ec1 in lipid bilayers within the detectable range of our measurements. This mutant appears to be more stable than H-ala, or the other helix-ala constructs, but is still significantly destabilized compared to the WT protein.

## DISCUSSION

In our study, we found that reducing non-polar side-chain interactions at the dimerization interface yields functional, monomeric CLC-ec1 in 2:1 POPE/POPG lipid bilayers. In our first approach, we found that an accumulation of 8-10 subtractive substitutions of phenylalanine, leucine and isoleucine to alanine per dimer, produce monomeric CLC-ec1 that behaves comparably to the monomeric I201W/I422W CLC-ec1 control (Chadda et al. 2016; Robertson, Kolmakova-Partensky, and Miller 2010). In our original hypothesis, we proposed that removal of these side-chains would lead to a formation of cavities at the dimerization interface that would selectively destabilize the dimer complex **(Figure 2A, C & Supplemental Figure 1A**). While our experimental findings agree with this hypothesis, we cannot be certain that cavities are being formed without high-resolution structural information about these constructs. For instance, the side-chains at the surface may re-pack, lipids may come in to form additional interactions, or the backbone may adjust to minimize this cavity volume and reclaim some of the protein interactions. One piece of evidence supporting the latter, is that H-ala shows a small decrease in functional activity, suggesting that backbone structural changes may occur. In addition, in our second approach, we find that a single mutant L194A, is also capable of shifting CLC-ec1 to the monomeric form in membranes. In examining the original interaction energy analysis, we find that L194 was the strongest favorable contributor to VDW packing between the two subunits (**Figure 1F**). Thus, while we do not know if these constructs possess the structural cavities in the dimer state as modeled, the behavior of the protein in lipid bilayers indicates that a loss of VDW interactions in the dimer state is correlated with decreases in dimerization.

While the behavior of the protein in lipid bilayers appears sensible, many questions arise when we consider the behavior of the protein in detergent. Surprisingly, several of the helix-ala constructs - I-ala, P-ala and Q-ala, purified as dimers in DM micelles. Thus, while we removed 8-10 side-chains at the dimerization interface, putatively leading to the formation of large cavities and a significant destabilization of the complex in membranes, we find that there is no indication of an effect in the DM micelle environment. In the following discussion, we will consider the several potential explanations for these contradictory observations.

First, it is possible that the protein may exhibit a stronger dimerization affinity in DM micelles compared to lipid bilayers. In this case, the protein is in a dynamic equilibrium, but the equilibrium is significantly shifted so that the dimer population is observed under the experimental conditions. However, despite observing dynamic equilibrium behavior for CLC-ec1 in 2:1 POPE/POPG lipid bilayers (Chadda et al. 2016), we observed no effect of the protein population due to dilution with detergent micelles. In addition, we found no evidence of subunit exchange when monitoring for an increase in heterodimer FRET signal when the protein was studied for up to a week. However, we did observe changes in L194A protein by SEC over a similar timescale, suggesting the protein population can evolve into monomers, but does not reassemble as dimers during the relaxation of the population. Thus, while a shifted equilibrium constant is still possible, our investigation indicates that the population does not participate in a dynamic equilibrium in the DM micelles over a week-long time-scale. In other words, the I-ala, P-ala and Q-ala dimers are kinetically trapped as dimers within the detergent micelles. This observation appears to be specific to the CLC-ec1 homologue as it is in contrast to the behavior that has been reported for CLC^F^-eca (Last and Miller 2015) or other proteins like Glycophorin-A (Fisher, Engelman, and Sturgis 1999, 2003). This is, however, in agreement with studies that describe how membrane proteins can exhibit high kinetic stability, as was demonstrated with the dissociation of diacylglycerol kinase trimers exhibiting a half-life of several weeks (Jefferson, Blois, and Bowie 2013). Note, this phenomenon may also be specific to the DM micelle condition, as DM is a detergent that is preferred for x-ray crystallography studies (Stetsenko and Guskov 2017). Altogether, these results indicate that the stoichiometry of a membrane protein in detergent micelles may not reflect the overall stability of the complex, but rather a kinetically trapped form of the protein obtained *en route* during the purification process.

This raises the question - when does the dimer form during purification of the protein? One possibility is that the process of protein synthesis and expression drives dimerization of the subunits in the *E. coli* membrane. Since little is understood about CLC-ec1 synthesis during over-expression conditions, the following is speculation. When over-expressing the protein in cells, it is possible that CLC-ec1 is synthesized and partitions into the membrane at a high local density, which drives the assembly of dimers in the membrane. In this case, even if the dimer equilibrium is significantly destabilized, the dimer forms may be captured in the detergent micelles provided the dissociation kinetics are slow. Alternatively, CLC-ec1 dimers might form co-translationally as a required intermediate along the protein folding pathway, again leading to the capture of dimers upon membrane extraction. Finally, it is possible that the environment of the *E. coli* inner membrane differentially drives CLC-ec1 dimerization due to changes in lipid composition, or the presence of other unknown cofactors that stabilize equilibrium dimerization in the helix-ala constructs. For example, while 2:1 POPE/POPG is an appropriate synthetic mimic of the overall *E. coli* polar lipid composition, we know that the biological membranes contain cardiolipin (Ames 1968), possess a large degree of variability in acyl chain composition (Kito et al. 1972; Schmidt et al. 2019) as well as other lipoidal molecules (Campagna, Miller, and Forman 2003; Robertson 2018). Other environmental factors, such as leaflet asymmetry, ionic conditions and pH could all contribute to shifting the reaction equilibrium in the cell, compared to what we observe in our 2:1 POPE/POPG membranes. Finally, it is possible that the protein exists as a monomer in the *E. coli* inner membranes, but during the step of detergent extraction, the protein to micelle ratio drives the over-filling of micelles. In this case, the experimental goals of maximizing protein yields while minimizing the amount of detergent, may unknowingly bias membrane proteins towards higher stoichiometries due to the random co-occupancy, a phenomenon known as forced co-habitation that has been shown to contaminate binding studies in detergent micelles (Kobus and Fleming 2005). Altogether, we recognize that there are many factors that may lead to the observation of CLC-ec1 dimers during purification, but none of these preclude our ability to conclude that removal of non-polar side-chains along the dimerization interface destabilize the dimer when in the equilibrium environment of the lipid bilayer.

The final surprising observation that arose during our study, is that while I-ala, P-ala and Q-ala form kinetically stable dimers in DM micelles, H-ala showed the unique behavior of converting to monomers rapidly after purification. Further investigation revealed that this effect depended on L194A, and this mutation was sufficient to convert CLC-ec1 to monomers in lipid bilayers (**Figure 5K**). Functional transport studies and x-ray crystallography demonstrate that L194A maintains function, fold and the ability to dimerize. How then does this single leucine to alanine mutation yield such striking changes in dimerization, in both detergent micelles and in lipid bilayers? Examination of the position of L194 shows that it is located at the symmetry interface of the inverted topology fold, which brings together the two halves of each subunit (Min et al. 2018). At this position, it is possible that L194 participates to stabilize the dimer, but also the dimerization interface, locking the H and P helices together at this seam. While L194A does not exhibit a change in function, H-ala showed a small reduction in transport activity, indicating that the H-P interaction is important for maintaining the functional fold of the dimerization interface. Other studies have indicated that structural changes may occur at the dimerization interface during the Cl^-^/H^+^ transport cycle. For example, cross-linking of D417C results in a decrease in Cl^-^ turnover rate by restricting the motions of the P-helix (Khantwal et al. 2016). In addition, a recent x-ray crystal structure of CLC-ec1 with mutations E113Q/E148Q/E203Q (PDB ID: 6V2J) yields a proposed outward-facing open state with a conformational change at the dimerization interface (Elvington, Liu, and Maduke 2009; Khantwal et al. 2016; Chavan et al. 2020). While we know that the dimer complex does not go through significant conformational changes due to the ‘straight-jacketing’ cross-linking experiments carried out previously at the Q and I helices (Nguitragool and Miller 2007), it is possible that localized changes do occur at the center of the dimerization interface during the transport cycle, particularly at the seam of the H-P helices. If these different conformational states disrupt the surface complementarity, then it is possible to observe modal stabilities for CLC-ec1 dimerization. Along these lines, we hypothesize that L194, in addition to being a potential hot-spot for VDW packing, participates in stabilizing the dimerization interface and locking in the high-affinity dimerization conformation. This hypothetical mechanism also proposes an increase in conformational dynamics at the dimerization interface, which may explain why L194A and H-ala exhibit the unique ability to break apart detergent micelles. Further studies examining possible structural changes, or subunit dynamics, will be necessary to understand why a single alanine mutation is capable of making CLC-ec1 a monomer in membranes.

## CONCLUSION

Here we report that CLC-ec1 dimerization in lipid bilayers is destabilized when non-polar side-chains are mutated to alanine at the membrane embedded dimerization interface. Despite this, we find that several constructs, where substantial side-chain contacts have been removed, can still be found as dimers upon purification in detergent micelles or during crystallization. In addition, we find that CLC-ec1 can be made monomeric with a single alanine substitution at residue L194, identifying a novel molecular hot-spot for dimerization. Altogether, these results demonstrate that interfacial side-chains impact dimer stability in lipid bilayers, and that membrane protein stoichiometry is a highly contextual observable that depends strongly on the solvent environment.

## ACKNOWLEDGEMENTS

The Robertson lab is supported by the National Institute of General Medical Science, National Institutes of Health (R01GM120260, R21GM126476). We are grateful to the staff at beamlines 5C at PALII (Pohang Light Source II, Pohang Accelerator Laboratory, Pohang, Republic of Korea) for assistance at the synchrotron. This work was partly supported by the KBRI Basic Research Program through Korea Brain Research Institute funded by the Ministry of Science and ICT, Korea (20-BR-01-05) to H.-H. L. We thank Rahul Chadda and the Robertson Laboratory for useful discussions.

## METHODS

### *In silico* van der Waals Energy Calculations

The atomic coordinates of the CLC-ec1 were taken from the x-ray crystal structure (PDB ID: 1OTS). The CHARMM-GUI (http://www.charmm-gui.org/) (Brooks et al. 2009; Lee et al. 2016; Jo et al. 2014; Jo et al. 2008) webserver was used to build missing hydrogens into the atomic coordinates and to mutate C85 to Ala. All sidechain atoms of the CLC-ec1 model structure were then energy-minimized for 8000 steps using NAMD (Phillips et al. 2005) and the CHARMM27 force-field. Pairwise van der Waals (VDW) interaction energies were calculated using NAMD energy between the sidechain of residue *i* on subunit *S* and the sidechain of residue *j* on subunit *S’*:

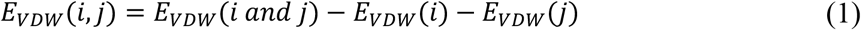

The total sidechain interaction energy term for the residue *i* on *S* was calculated as the summation of all pairwise interaction energies between the sidechain of this residue and the sidechains of all residues on the opposing subunit *S’*:

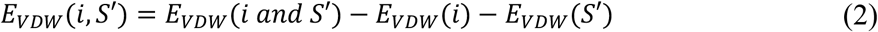

The van der Waals interactions were truncated at the cutoff distance of 12 Å and the interaction potential was truncated by a smooth switching function at the cutoff distance of 8.0 Å. Otherwise, default parameters were used.

### Cavity Volume Calculations

The CHARMM-GUI webserver was used to build missing hydrogen atoms into the WT CLC-ec1 1OTS structure, and generate the multiple alanine substitutions for H-ala, I-ala, Q-ala and P-ala. The cavity volumes at the dimerization interface were calculated using two approaches. First, we used *trj_cavity* 2.0 with a 1.4 Å grid spacing (Paramo et al. 2014). The difference in cavity volumes were calculated for each of the helix-ala structures compared to WT. The second method used the webserver CASTp 3.0 (http://sts.bioe.uic.edu/castp/index.html?2cpk) (Tian et al. 2018), defining the boundaries of the protein by its molecular surface and using 1.4 Å radius sphere. Spheres showing the cavity volumes located at the sites of the alanine substitutions are the representative negative spaces generated by the CASTp webserver when performing the calculations.

### CLC-ec1 constructs

Wildtype (WT) CLC-ec1 was cloned into a pASK90 vector encoding a hexa-histidine tag at the C-terminus. All mutations were made by Quickchange II Site-Directed Mutagenesis (Agilent, Santa Clara, CA) followed by DNA sequencing of the full gene. ‘WT’ CLC-ec1 contains two substitutions, C85A and H234C, to allow for site-specific labeling on each subunit. The molecular weights (MW) of each construct are as follows: MW_WT(C85A/H24C)_ = 51997 g/mol, MW_H-ala(C85A/H234C/L194A/I197A/L198A/I201A_ = 51829 g/mol, MW_I-ala(C85A/H234C/F219A/I220A/I223A/I227A_ = 51795 g/mol, MW_P-ala(C85A/H234C/L406A/I409A/I410A/L413A)_ = 51829 g/mol and MW_Q-ala (C85A/H234C/I422A/L423A/I426A/L430A/L434A)_ = 51829 g/mol. Each construct has an extinction coefficient *ε* = 46020 M^−1^ cm^−1^. Molecular weights and extinction coefficient were calculated using the Peptide Property Calculator (http://biotools.nubic.northwestern.edu/proteincalc.html).

### Solutions

Breaking buffer (BB): 100 mM Sodium Chloride (NaCl, Research Products International (RPI), Mount Prospect, IL), 50 mM Tris (RPI), 5 mM Tris(2-carboxyethyl)phosphine HCl (TCEP, Soltec Ventures, Beverly, MA), pH 7.5. Cobalt wash buffer (CoWB): 100 mM NaCl, 20 mM Tris, 1 mM TCEP, 5 mM n-Decyl-B-D-Maltopyranoside (DM, Anatrace, Maumee, OH), pH 7.5. Size exclusion buffer (SEB): 150 mM NaCl, 20 mM 3-(N-morpholino)propanesulfonic acid (MOPS, RPI), 5 mM DM, pH 7.0. Dialysis buffer (DB): 300 mM Potassium Chloride (KCl, RPI), 20 mM Citrate (RPI), pH 4.5. External Buffer (EB): 150 mM Potassium Sulphate (K2SO4, Sigma-Aldrich, Saint Louis, MO), 1 mM KCl, 20 mM Citrate, pH 4.5.

### Protein Purification

Protein purifications were carried out as described previously (Chadda et al. 2016; Robertson, Kolmakova-Partensky, and Miller 2010; Maduke, Pheasant, and Miller 1999; Walden et al. 2007; Accardi and Miller 2004). Competent BL21-AI *E. coli* cells (Invitrogen, Carlsbad, CA) were transformed with a plasmid (pASK90 vector) that expresses the respective WT CLC-ec1 and CLC-ec1 alanine variants. Two flasks of 1 L volumes of Terrific Broth (RPI) supplemented with 4 mL of glycerol each, were inoculated with the transformed *E. coli* in the presence of 100 μg/mL ampicillin (GoldBio, Olivette, MO). The cultures were incubated at 37 °C, 220 RPM. Protein expression was induced when the growth reached OD_600_ = 1.0 with 0.2 μg/mL anhydro-tetracycline in dimethylformamide (Sigma-Aldrich). The cultures were incubated for an additional 3 hours before harvesting (5020 g, 20 minutes, 4°C) and the pellet was stored overnight at 4°C. The next morning, cell pellets were resuspended in BB to a volume of 50 mL with 10 μg lysozyme (RPI), 10 μg DNAase (Sigma-Aldrirch), 1 mM PMSF (RPI), 0.6 μM aprotinin (GoldBio), 0.84 μM leupeptin (Thermo Fisher Scientific, Waltham, MA), 0.56 μM pepstatin (Thermo Fisher Scientific) and lysed by sonication. Membrane proteins were extracted with 2% (*w/v*) DM while rotating at room temperature for 2 hours. Cell debris was pelleted down (30,966 g, 45 minutes, 4 °C), and the supernatant was loaded onto a cobalt metal affinity column (Takara Bio, Kusatsu, Japan) equilibrated in CoWB. The column was washed with CoWB containing 20 mM Imidazole (RPI), and protein eluted with CoWB containing 400 mM Imidazole (RPI). The eluate was concentrated with a 10,000 MWCO centrifugal filter (Millipore-Sigma, Saint Louis, MO), which was then injected onto a Superdex 200 Increase size exclusion column (GE Life Sciences, Little Chalfont, UK) equilibrated in SEB using a Biorad NGC System. Protein samples were collected using a fraction collector and then quantified by absorbance at a wavelength of 280 nm using a Nanodrop 2000c UV-visible light spectrophotometer (Thermo Fisher Scientific).

### Cysteine Accessibility Assay

Cysteine accessibility assays were carried out as described previously (Chadda and Robertson 2016). WT CLC-ec1 and helix-ala constructs have an alanine substitution at C85 and a cysteine substitution at H234. C85A/H234C CLC-ec1 have a single solvent accessible cysteine while in DM micelles, and two buried endogenous cysteines, C302 and C347 that become reactive when the protein is denatured with SDS (RPI). To measure cysteine accessibility, Ellman’s reagent or 5,5’-Dithio-bis(2-nitrobenzoic acid) (DNTB, Sigma-Aldrich) is added to the protein, whereupon DNTB reacts with the thiolate form of the cysteine in a disulfide exchange reaction to yield a stoichiometric product of TNB^2-^ dianion that absorbs light at 412 nm. In the assay, a 10 mM stock of is freshly prepared in 100 mM Sodium Phosphate (RPI), 1 mM EDTA (Macron, Center Valley, PA), pH 8.0. Before the measurement, a 1:1 dilution of Ellman’s stock is prepared with SEB to make a 5 mM Ellman’s reagent working solution. A 10 μM protein sample is prepared in SEB at a volume of 300 μL, and loaded into a 1 cm quartz cuvette in a Nanodrop 2000c UV-VIS spectrophotometer, where the absorbance at 412 and 750 nm is measured every 30 seconds. After two minutes, 20 μL of the working Ellman’s reagent solution is added to the cuvette and the sample is mixed. The absorbance is followed for 15 minutes, after which 40 μL of 20% SDS (*w/v*) in 150 mM NaCl, 20 mM MOPS, pH 7.0 is added and mixed. The sample is monitored for an additional 30-40 minutes until the absorbance at 412 nm stabilizes. The final TNB^2-^ absorbance is corrected for baseline shifts by calculating A_412_ - A_750_. These values were additionally corrected by subtracting for the intrinsic background from Ellman’s reagent, A_412,Ellman’s,DM_ = 0.0684 and A_412,Ellman’s,SDS_ = 0.0631, measured by carrying out the assay in the absence of protein. The concentration of [TNB^2-^] is calculated from the data that has been baseline and background corrected using the following extinction coefficients: ε = 1.38 x 10^−8^ M^−1^ cm^−1^ in DM buffer and ε = 1.24 x 10^−8^ cm^−1^ M^−1^ in SDS buffer. Data was fit with a single exponential association function to quantify the plateau absorbance for each reaction condition.

### Analytical size exclusion chromatography

Immediately after initial purification, the protein samples were diluted to a protein concentration of 10 μM in SEB containing 5 mM DM. This yielded a controlled protein to detergent mole fraction of χ = 3 x 10^−3^ subunits/detergent molecule. The protein sample was then incubated at 4 °C for the duration of the experiment. At various time points, 400-450 μL of the sample was filtered and then injected on to a Superdex 200 Increase 10/300 GL size exclusion column equilibrated with SEB. The chromatography spectrum measured at 280 nm was baseline corrected and area normalized between 10-16 mL elution volume. The curve was fit to a sum of two Gaussian functions, and the fraction of dimer was quantified as the fractional area under the dimer peak. For the DM dilution studies of P-ala, we diluted the protein to a concentration of 20 μM in SEB containing 5 mM DM for a final mole fraction of χ = 6 x 10^−3^ subunits/detergent immediately after the initial purification. Then, P-ala was diluted 1:1 with SEB containing either 100 mM or 1 M DM, for final protein concentrations of 10 μM and final DM concentrations of 52.5 mM and 502.5 mM. This corresponds to mole fraction values of χ = 2 x 10^−4^ subunits/detergent and χ = 2 x 10^−5^ subunits/detergent, respectively. After increasing the amount of detergent in the P-ala samples we immediately placed the samples at 4°C and incubated the samples for six days. After the six day incubation period, 400-450 μL of the sample was filtered and then injected on to a Superdex 200 Increase size exclusion column equilibrated with SEB.

The CLC-ec1 L194A C85A samples were concentrated by adding a volume of protein sample at 10-20 μM (χ = 3-6 x 10^−3^ subunits/detergent) to cobalt metal affinity beads, Column Volume (CV) = 100 μL in a Micro Bio-Spin Chromatography Column (Bio-Rad, Hercules, California) equilibrated in SEB buffer and eluting the sample in a smaller volume of SEB containing 400 mM Imidazole. Concentrated protein was quantified by a Bradford Assay (Bio-Rad). Dilution of the CLC-ec1 L194A C85A constructs were either carried out by adding more SEB to the sample, i.e., diluting the protein while keeping the detergent concentration constant. We also diluted the samples by increasing the amount of detergent by adding SEB with a concentration of DM detergent at 100 mM to the sample or by completing the size exclusion chromatography step at the end of the initial purification with the size exclusion column equilibrated in SEB containing 20 mM DM. After concentrating or diluting the samples 400-450 μL of sample was filtered and injected on to the size exclusion column. After some time, 1-6 days at an incubation temperature of 4 °C, another 400-450 μL of sample was filtered and injected on to the size exclusion column.

### Site specific Cy3 and/or Cy5-maleimide labeling of CLC-ec1

Fluorophore-maleimide labeling of C85A/H234C CLC-ec1 and the helix-ala constructs variants were carried out as described previously (Chadda and Robertson 2016). For the single-molecule total internal reflection fluorescence (TIRF) microscopy studies, protein at a concentration of 10 μM in SEB was reacted with 50 μM Cy5-maleimide (Lumiprobe, Hunt Valley, MD) for 15 minutes before the reaction was quenched with 5 mM cysteine (RPI). The reacted protein was batch bound to cobalt metal affinity beads, CV = 100 μL (Takara Bio) in a Micro Bio-Spin Chromatography Column equilibrated with CoWB. Excess fluorophore was washed away with 5 CV of CoWB, before eluting the labeled protein with CoWB containing 400 mM imidazole (RPI). To remove the imidazole, which interferes with the protein quantification, the sample was loaded onto a freshly poured 3 mL G50 Sephadex column (GE Life Sciences) equilibrated in CoWB, and the visible eluant was collected drop by drop. The labeling yield was quantified by placing the sample in a 1 cm quartz cuvette in a Nanodrop 2000c UV-VIS spectrophotometer and measuring the absorbance spectrum between 190-840 nm. The protein concentration in the presence of Cy5, [*protein · Cy5*], is calculated as (eqn. 3):

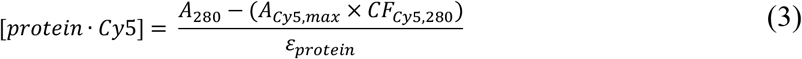

where *A*_280_ is the absorbance at 280 nm, *A_Cy5,max_* is the peak absorbance of Cy5 ≈ 649 nm, *CF_Cy5,280_* = 0.02, is the correction factor for the absorbance of Cy5 at 280 nm, and *ε_protein_* is the extinction coefficient for the protein at 280 nm. The subunit labeling yield, *P_Cy5_*, is calculated as (eqn. 4):

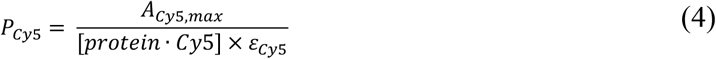

where, *ε_Cy5_* = 2.5 x 10^5^ M^−1^ cm^−1^ is the extinction coefficient for Cy5 at 649 nm. For the Förster Resonance Energy Transfer (FRET) studies, protein was labelled with Cy3-maleimide (Lumiprobe) or simultaneously co-labelled with Cy3- and Cy5-maleimide. For Cy3-labeling alone, the same procedure was followed, except that the quantification is corrected for Cy3 contribution at 280 nm. Thus, the protein concentration in the presence of Cy3, [*protein · Cy3]*, is calculated as (eqn. 5):

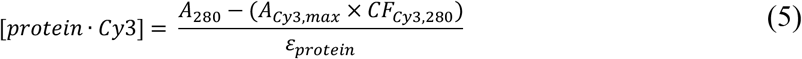

where *A_Cy3,max_* is the peak absorbance of Cy3 ≈ 552 nm, and *CF_Cy3,280_* = 0.08 is the correction factor for the absorbance of Cy3 at 280 nm. The subunit labeling yield, *P_Cy3_*, is calculated as (eqn. 6):

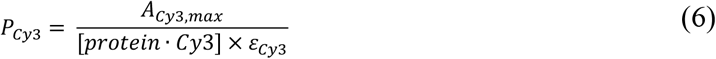

where, *ε_Cy3_* = 1.5 x 10^5^ M^−1^ cm^−1^ is the extinction coefficient for Cy3 at 552 nm. For labeling of Cy3 and Cy5 simultaneously, there are two correction factors to consider, Cy3 and Cy5 absorbance at 280, as well as the contribution of Cy5 absorbance in the Cy3 peak. Thus, the protein concentration in the presence of Cy3 and Cy5, [*protein · Cy3/Cy5]*, is calculated as (eqn. 7):

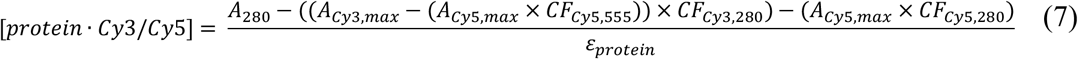

where *CF_Cy3,565_* = 0.08 is the correction factor for the absorbance of Cy5 around the Cy3 peak. The subunit labeling yield of Cy3 in the presence of Cy5, *P_Cy3:Cy5_*, is calculated as (eqn. 8):

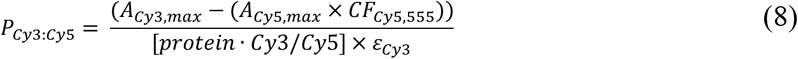

and the subunit labeling yield of Cy5 in the presence of Cy3, *P_Cy5:Cy3_*, is calculated as (eqn. 9):

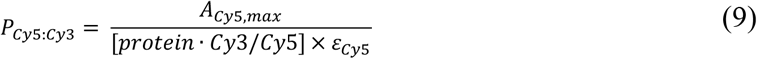

### Bulk FRET measurements of CLC-ec1 in detergent

For bulk FRET measurements to test for heterodimer formation indicative of subunit exchange, samples were prepared as Cy3-labeled, Cy5-labeled, “co-labeled” Cy3/Cy5-labeled protein that involved simultaneous labeling with both fluorophores, or “mixed” samples consisting of Cy3-labeled protein and Cy5-labeled protein added to the same well. Protein samples were studied at 1 μM in SEB. Samples were incubated at the indicated temperature and measured in a 96-well half area plate (Corning, Corning, NY) sealed with a film cover between experiments (Excel Scientific, Victorville, CA). Fluorescence intensity measurements were made using a TECAN Sapphire II plate reader. Donor spectra were collected by exciting the samples at 540 nm and collecting emission spectra between 555-850 nm. Acceptor spectra were collected by exciting samples at 640 nm and collecting the emission spectra between 653-850 nm. The direct excitation of Cy5 was subtracted away from the donor spectra of the experimental sample using the donor spectra from a sample labeled with only Cy5 (eqn. 10) (Granier et al. 2007).

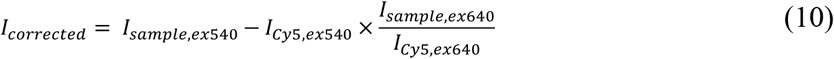

The corrected spectra were area normalized and the FRET efficiency, *E*, was calculated by donor quenching (eqn. 11):

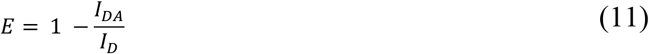

where *I_DA_* is the fluorescence intensity of the donor (Cy3) in the presence of the acceptor (Cy5) and *I_D_* is the fluorescence intensity of Cy3 in the absence of the Cy5.

### X-ray crystallography

Expression and purification of the C85A/L194A and C85A/L194A/H234C mutant CLC-ec1 proteins were performed as described above. Briefly, C-terminal His_6_-tagged CLC-ec1 cloned in the pASK vector was transformed in *E.coli* BL21(DE3) (Invitrogen, Carlsbad, CA) and the expression of CLC-ec1 protein was induced with anhydrotetracycline (Acros Organics, 0.2 μg/mL) for 3 hrs at 37°C. The mutant CLC-ec1 proteins were extracted with n-decyl-β-D-maltopyranoside (DM, Anatrace; 2% (*w/v*)) and allowed to bind to cobalt resin (Talon, Clontech; 1 mL bed volume/L culture). The column was washed sequentially with cobalt wash buffer (CoWB; 100 mM NaCl, 20 mM Tris-HCl_4_, and 5 mM DM, pH 7.5) and CoWB containing 20 mM imidazole until UV (OD_280_) signals reached the baseline. Proteins were eluted with CoWB containing 400 mM imidazole and concentrated to 5-10 mg/mL by using Amicon ultra-50K (Millpore-Sigma). For crystallizing mutant CLC-ec1 proteins, the C-terminal His-tags were removed by endoproteinase LysC (New England Biolab, 0.5 μg/mg protein, Ipswich, MA) and each mutant protein was complexed with F_AB_ fragment (1:1.6, CLC-ec1:Fab fragments (OD_280_ ratio)) at room temperature for 30 min. (Dutzler, Campbell, and MacKinnon 2003). The complexes were purified on a Superdex 200 increase size-exclusion column (GE healthcare) equilibrated with 100 mM NaCl, 10 mM Tris-HCl, and 5 mM DM, pH 7.5, concentrated to 10-20 mg/mL, and mixed with an equal volume of crystallization solution in a hanging-drop vapor-diffusion chamber.

The C85A/L194A and C85A/L194A/H234C mutant CLC-ec1 protein crystals were grown in 29% (w/v) PEG400 (Sigma-Aldrich), 200 mM NaCl (Sigma-Aldrich), 100 mM Hepes-NaOH(Sigma-Aldrich), pH 7.0, and 25% (w/v) PEG400 (Sigma-Aldrich), 100 mM NiNO_3_ (Sigma-Aldrich), 100 mM Glycine (Sigma-Aldrich), pH 9.5, respectively, in 3~10 days at 22 °C. Cryo-protection was performed by the slow increases of PEG concentration in the mother liquor to ≈ 35% before freezing crystals in liquid nitrogen. X-ray datasets were collected on beamline BL-5C at the Pohang Accelerator Laboratory II (PAL II, Pohang, Korea) at an X-ray wavelength of 1 Å. Data were processed and scaled by using HKL2000, and initial models were generated by molecular replacement using Phaser (McCoy et al. 2007; McCoy 2007) in the CCP4 software suite (Winn et al. 2011) against a wildtype CLC-ec1 F_AB_ complex, 4ENE (Lim, Shane, and Miller 2012). Rigid-body refinements were performed in REFMAC5 (Murshudov, Vagin, and Dodson 1997) and the models were further refined in Phenix ver. 1.13 (Liebschner et al. 2019). Final models were examined and manually adjusted by using the *Coot* software (Emsley and Cowtan 2004). Structures are uploaded on the PDB as 7CVT for the C85A/H234C/L194A and 7CVS for the C85A/L194A mutants. Crystallographic statistics are reported in **Supplementary Table 3**.

### Reconstitution of Protein into Liposomes

Reconstitution of WT CLC-ec1 and the alanine substituted constructs were carried out as described previously (Chadda et al. 2018). Briefly, 1-palmitoyl-2-oleoyl-sn-glycero-3-phosphoethanolamine (POPE, Avanti Polar Lipids Inc., Alabaster, AL) and 1-palmitoyl-2-oleoyl-sn-glycero-3-phospho- (1’-rac-glycerol) (POPG, Avanti Polar Lipids Inc.) in 25 mg/mL chloroform stocks, were mixed together in a glass vial in a ratio of 2 parts POPE to 1 part POPG (*v/v*). The lipids were dried by evaporation of the chloroform under a stream of nitrogen gas until a thin dried film of lipids accumulated at the bottom and sides of the vial. The film was washed with pentane (Sigma-Aldrich), before repeating the evaporation process 1-2 more times. Then, lipids were resolubilized in DB with 35 mM CHAPS (Anatrace) for a final concentration of 20 mg/mL lipids. The lipid-detergent mixture was solubilized using a cup-horn sonicator (Qsonica, Newtown, CT) until the sample was transparent. Protein was added to the solubilized lipid-detergent mixture and placed into 10,000 MWCO dialysis cassettes (Thermo Fisher Scientific), and then the samples were then dialyzed in the dark at 4 °C against 4 L of DB, with buffer changes every 8-12 hours for a total of 4 changes. At the end of dialysis, the samples were homogenized and extracted from the cassettes. As a final step, the samples were freeze-thawed to form large multi-lamellar vesicles (MLVs). This involved 7 repetitions of freezing the samples in either a dry ice ethanol mixture or liquid nitrogen and thawed in a room temperature water bath. The resultant MLVs are cloudy and settle to the bottom of the tube over time. Samples were stored at room temperature, in the dark for the desired amount of time, supplemented with 0.02% sodium azide (Sigma-Aldrich) to prevent contamination over time.

### Chloride Transport Assay

Chloride transport assays were carried out as described previously (Walden et al. 2007; Chadda et al. 2016). Proteoliposomes were prepared as described above, at a density of 1 μg/mg protein per lipid, corresponding to a reconstituted mole fraction of *χ_reconst_*. = 1.4 x 10^−5^ subunit/lipid. Freeze-thawed MLVs were extruded 21 times through a 0.4 μm polycarbonate membrane (Whatman, Maidstone, UK) using a LiposoFast-Basic extruder (Avestin, Ottawa, Canada). Immediately before the measurement, the external buffer of the proteoliposome sample was exchanged using a column of G50 Sephadex (GE Life Sciences) equilibrated in EB. The chloride efflux from the liposomes was monitored in EB using a homebuilt silver chloride electrode or a chloride ion sensing electrode (Cole-Parmer, Vernon Hills, IL). In general, the assay was conducted as follows. 10 μL of 10 mM KCl was added to 1.8 mL of EB in the recording chamber, followed by the addition of the proteoliposome sample in EB (100 μL extruded sample diluted to ≈ 200 μL volume after exchange). Transport was initiated by the addition of 1 μM valinomycin (Sigma-Aldrich) and 2 μM FCCP (Sigma-Aldrich) dissolved in methanol. To measure the total amount of intraliposomal chloride, 40 μL of 1.5 M n-Octyl-β-D-Glucopyranoside (β-OG, Anatrace) was added at the end of each transport experiment to release the remaining trapped chloride from inactive liposomes. Measurements made using the ion sensing electrode were carried out in the same manner with the exception that the starting volume of the reaction chamber was 3.8 mL, and 50 μL of 1.5 M β-OG was added at the end of each run. The time dependent transport data was fit to a two-component exponential relaxation function to determine the rate of chloride efflux, *k_CLC_*, and the fraction of trapped chloride, *F_Cl,0_*, after normalizing the data for the amount of total chloride (eqn. 12):

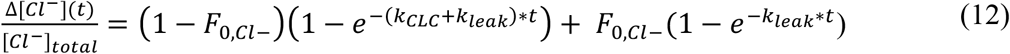

The rate of chloride leaking from empty 2:1 POPE/POPG liposomes, *k_Leak_*, was set as a constant of, 0.008 rel. Cl^-^/ sec, based on previous measurements (Chadda et al. 2016).

### Single-molecule photobleaching analysis of CLC-ec1 in lipid bilayers

Photobleaching experiments were carried out as described previously (Chadda et al. 2018; Chadda et al. 2016). Imaging was performed on an objective based total internal reflection fluorescence (TIRF) microscope, set-up for the imaging of single molecules, constructed by the Robertson Laboratory (Chadda et al. 2016). The microscope is equipped with a 637 nm OBIS laser (Coherent, Santa Clara, Coherent) that was used to excite Cy5 fluorophores on the protein. The laser intensity was adjusted to be between 150-275 μW, to maximize the signal to noise and obtain long photobleaching traces. Glass slides (Gold-Seal 24 x 60 mm no. 1.5 thickness, Thermo Fisher Scientific) and Coverslips (25 x 25 mm no. 1.0 thickness, Thermo Fisher Scientific) used for imaging were cleaned by placing five slides/coverslips into plastic slide holders at once. The glass was cleaned by filling the slide holders with 0.1% Micro-90 detergent (Cole-Parmer) and sonicating for 30 minutes before rinsing the slides fifteen times with de-ionized water. Then glass was cleaned by filling the slide holders with 100% ethanol (Thermo Fisher Scientific) and sonicating for 30 minutes before rinsing the slides fifteen times with de-ionized water. Finally, the slides were sonicated for 5 minutes in 0.2 M KOH (Sigma-Aldrich) followed by rinsing the slides fifteen times with de-ionized water before storing the slides in de-ionized water within a laminar flow cabinet. Silicon grease (Dow Corning, Midland, MI) was applied to the cleaned coverslips to create a flow-cell for imaging. After loading the slide onto the microscope, 35 μL of sample was loaded onto a flow cell and then the channel was washed multiple times with filtered DB to remove any excess proteoliposomes not bound to the glass. The number of spots corresponding to proteoliposomes with labeled proteins was kept at a low density, less than 600 spots in the imaging field, by diluting the samples with DB if needed. The average number of spots in each field imaged was 198 ± 17 (mean ± sem) and the total number of spots imaged for each measurement was 426 ± 9 (mean ± sem). Before imaging, the sample was passed through a 0.4 μM polycarbonate membrane (Whatman) using a LiposoFast-Basic extruder. All imaging was carried out in DB, which had been filtered through a 0.22 μM filter (Millipore-Sigma).

The analysis of the photobleaching traces was carried out as described previously (Chadda et al. 2018; Chadda et al. 2016). Image files were analyzed using a MATLAB-based CoSMoS analysis program (Friedman and Gelles 2015). Fluorescent spots were automatically selected by the image analysis software on the criteria of intensity. Selected spots selected were analyzed using a 4 x 4 pixel area of interest (AOI) centered around the peak. The total pixel intensity within each AOI was integrated as a function of time, and then the intensity traces were examined for step-like decreases in intensity indicating irreversible photobleaching of fluorophores.

### Quantifying *F_Dimer_*

The calculation of the amount of dimer, *F_Dimer_*, for the CLC-ec1 multiple alanine variants was carried out as described previously (eqn. 14) (Chadda et al. 2016). The experimental photobleaching data 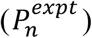 was fit to a monomer control distribution 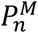, determined from measurements of I201W/I422W, a constitutive CLC-ec1 monomer (Robertson, Kolmakova-Partensky, and Miller 2010; Chadda et al. 2018), the dimer control distribution 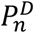, determined from R230C/L249C CLC-ec1 (Nguitragool and Miller 2007; Chadda et al. 2018), a constitutive CLC-ec1 dimer, by least-squares analysis of the residuals, *R^2^*.

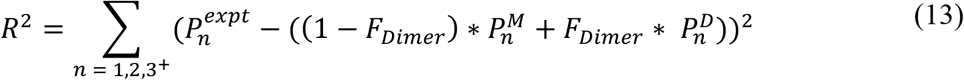

to estimate the best-fit of *F_Dimer_* as a function of χ*. From there, F_Dimer_ vs. χ* is fit to an equilibrium dimerization isotherm:

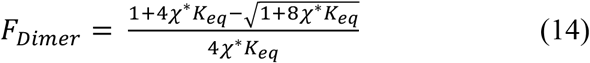

for estimation of the binding curve and the lower limits on the equilibrium constant for the reaction.

## SUPPLEMENTARY FIGURES & TABLES

**Supplemental Figure 1.**
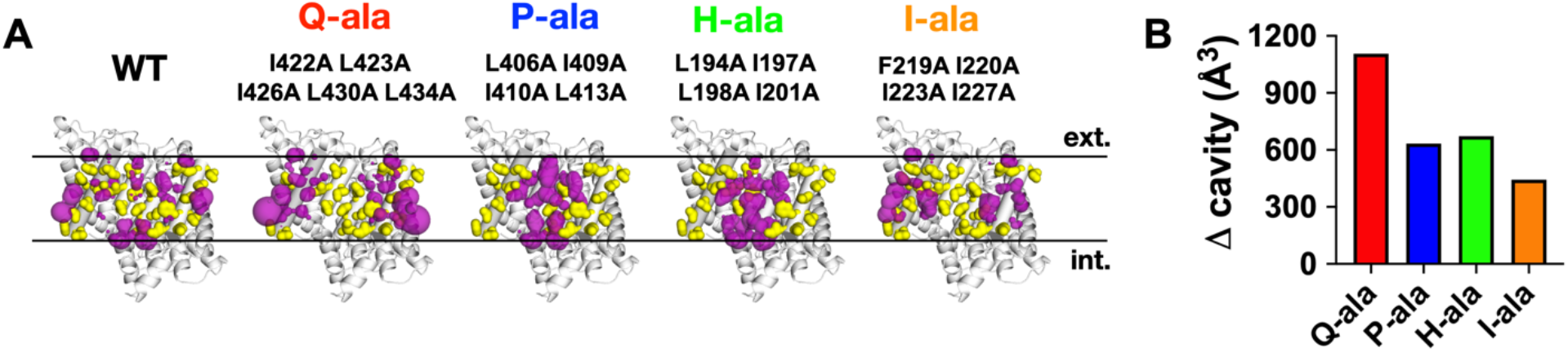
Cavities form in multiple alanine variants due to subtractive substitutions to alanine. (A) Coordinates of a CLC-ec1 structure (PDB ID: 1OTS), were manipulated in the webserver CHARMM-GUI to model substitutions from the native residue to alanine (substitutions are listed above each computationally created structure) and to place hydrogens into each structure. Analysis in the webserver CASTp found every cavity within each structure. The cavities that are close to the dimerization interface in the WT structure are shown here in magenta. While the cavities that were either created or grew larger due to the alanine substitutions are shown in each helix-ala variant. (B) Data here show changes in cavity volumes of the multiple alanine variants in comparison to the WT CLC-ec1 structure (the sum of a multiple alanine variant’s cavity volumes - the sum of the WT cavity volumes).

**Supplemental Figure 2.**
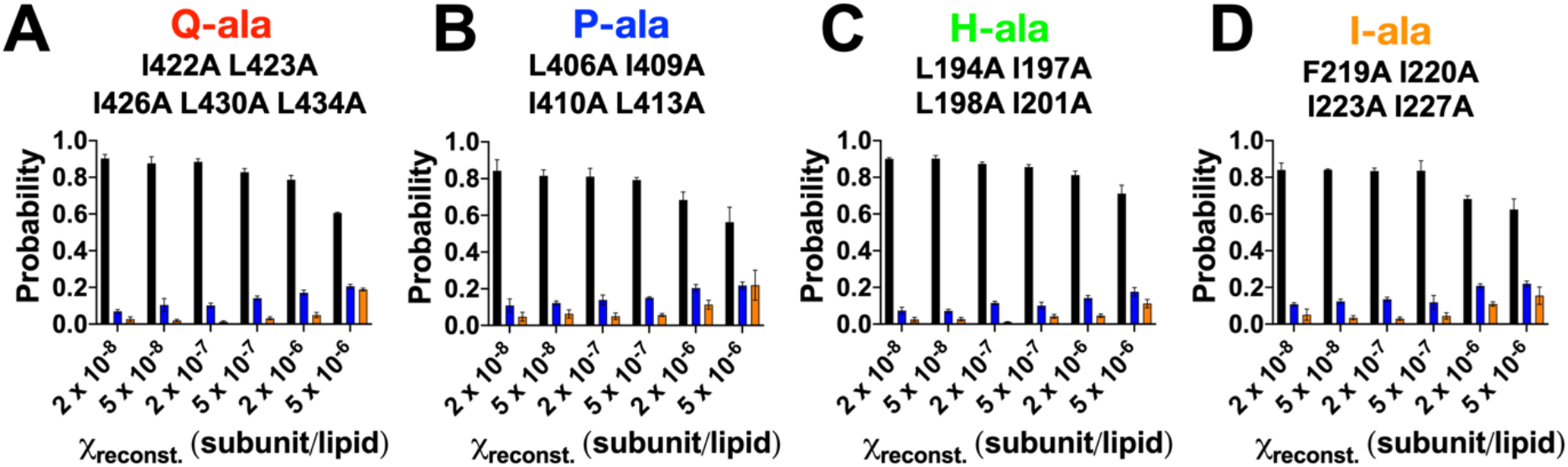
Q-ala, P-ala, H-ala, and I-ala are monomeric in 2:1 POPE/POPG membranes. (A, B, C, D) Photobleaching step distribution (black - one step photobleaching events, blue - two step photobleaching events, and orange - three or more step photobleaching events) mean ± sem, n = 3.

**Supplemental Figure 3.**
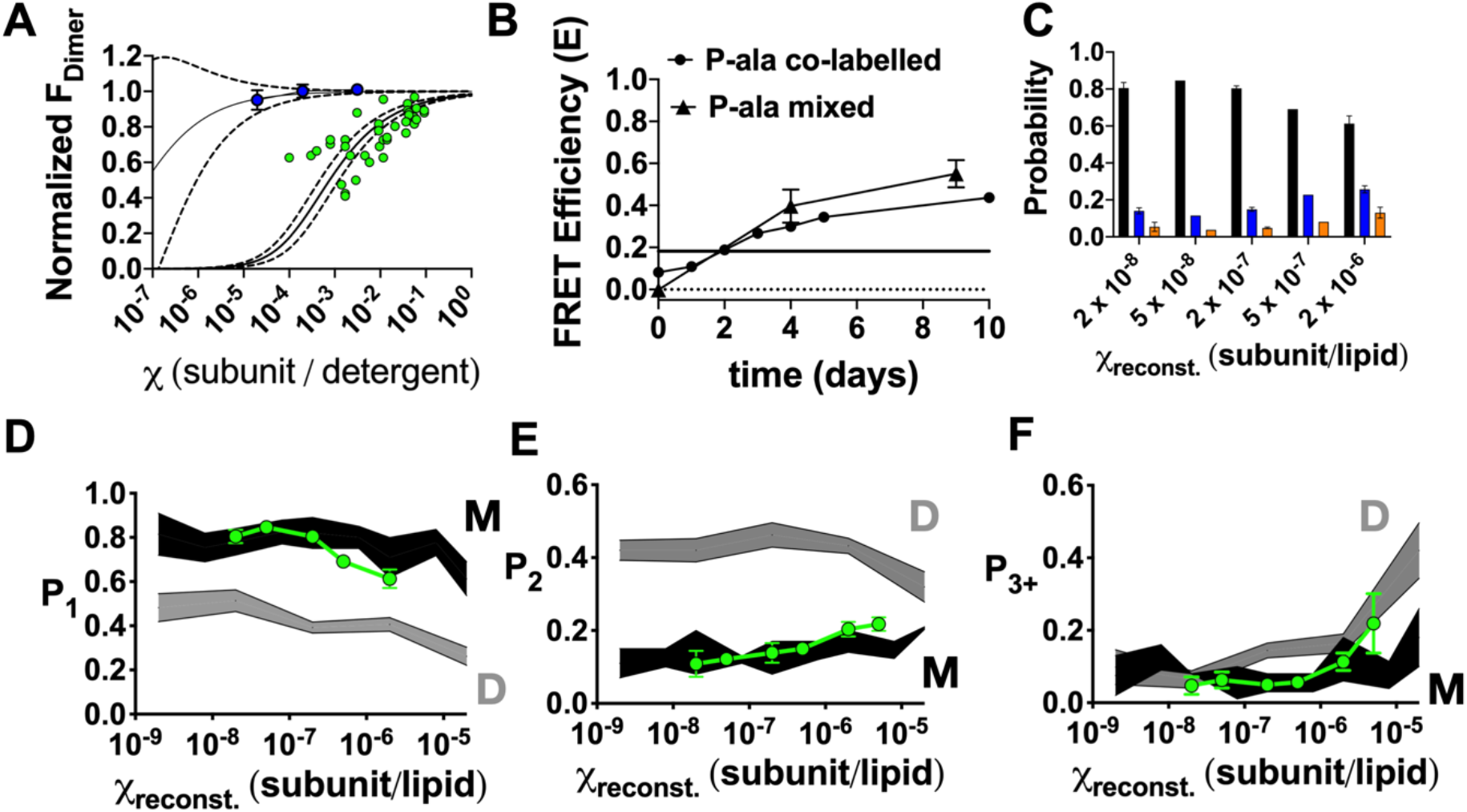
(A) Dilution studies of P-ala (blue) and L194A (green). Samples were either concentrated or diluted (see methods) and run on a size exclusion column. The data was fit with the sum of two Gaussians to calculate the fraction of dimer, F_Dimer_. The F_Dimer_ values were then normalized to the CLC-ec1 that exists as a monomer in detergent and lipids “I422W/I201W” data and the WT CLC-ec1 data which exists as a dimer in detergent micelles to calculate the Normalized F_Dimer_ values. For each construct the data set was fit with a dimerization isotherm (solid black line) and the dotted lines representing 95% CI. P-ala data each circle is the mean ± sem while each measurement made with the L194A data set is shown. (B) FRET analysis of the P-ala-Cy3 and P-ala-Cy5 subunit-exchange experiment in 5 mM DM, showing a change of the FRET signal as a function of protein incubation at 37°C. FRET represents donor quenched bulk FRET, *E* = 1-*F_AD_/F_D_*, ‘mixed’ refers to combining protein P-ala-Cy3 and P-ala-Cy5 after separate labeling, whereas ‘co-labelled’ refers to the positive control of P-ala that was labelled with Cy3 and Cy5 simultaneously. Experiments were carried out in SEB. As co-labelled controls are the control for a fully mixed sample, the development of FRET in these samples indicates that an aggregation process is occurring. The plateau of the P-ala co-labelled samples incubated at RT is represented as a solid black line at 0.18. Co-labelled P-ala sample n = 2 (acceptor/donor mole fraction = 0.22 ± 0.0) and P-ala mixed samples n = 3 (acceptor/donor mole fraction = 0.8). (C) Photobleaching step distribution (black - one step photobleaching events, blue - two step photobleaching events, and orange - three or more step photobleaching events) mean ± sem, n = 1-5. (D) *P_1_*, (E) *P_2_*, and (F) *P_3+_* photobleaching probabilities as a function of the reconstituted mole fraction, *χ_reconst_*. (subunits/lipid). Data represent mean ± sem, n = 1-5. CLC-ec1 monomer (M – black, I201W/I422W) and dimer (D – grey, R230C/L294C) control data (Chadda et al., 2018) are shown for reference (mean ± standard error bands). Experiments were carried out in dialysis buffer, 300 mM KCl, 20 mM citrate, pH 4.5.

**Supplementary Table 1.**
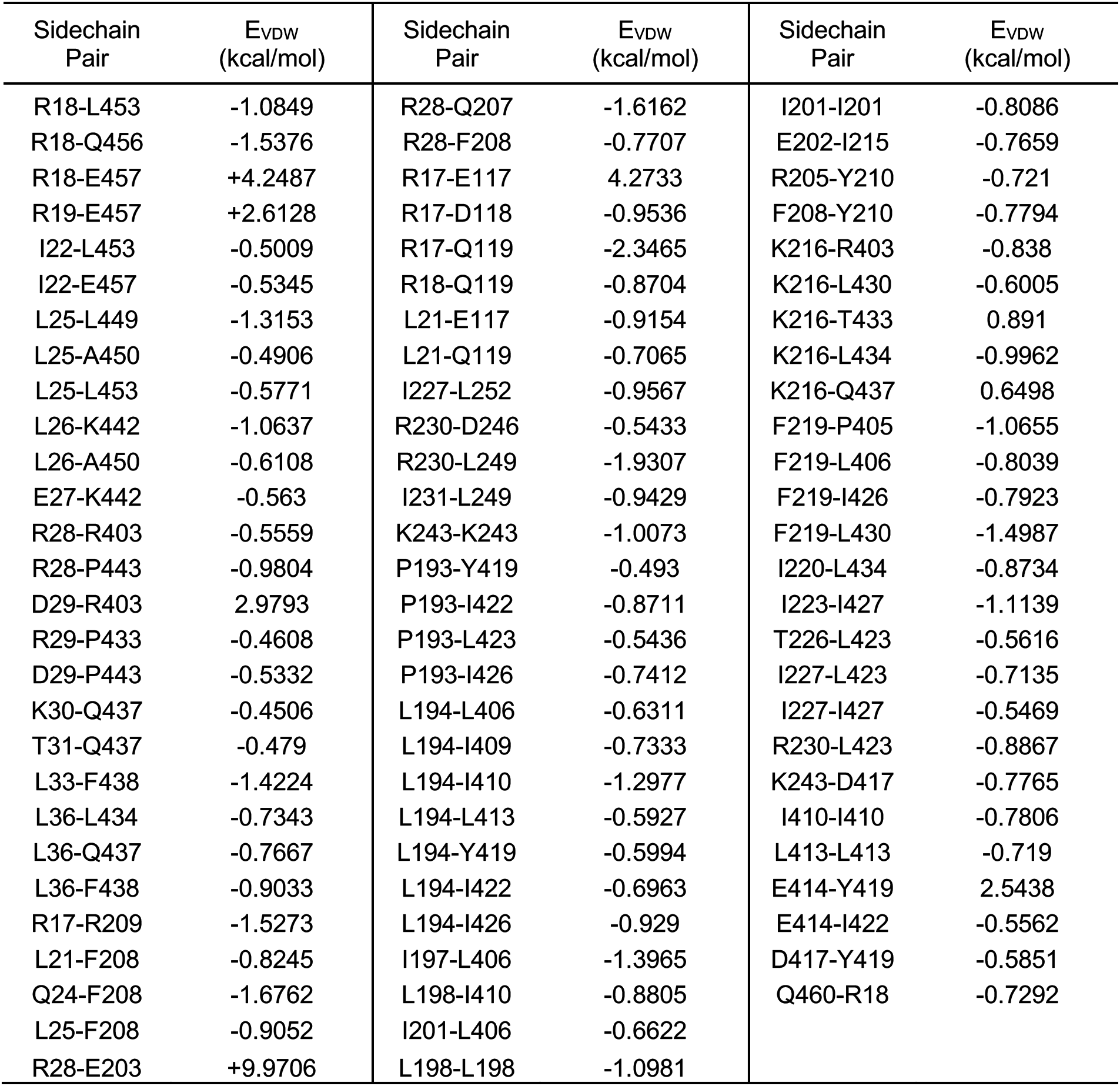
Pairwise side-chain VDW interaction energies.

| Sidechain Pair | E<sub>VDW</sub> (kcal/mol) | Sidechain Pair | E<sub>VDW</sub> (kcal/mol) | Sidechain Pair | E<sub>VDW</sub> (kcal/mol) |
| --- | --- | --- | --- | --- | --- |
| R18-L453 | -1.0849 | R28-Q207 | -1.6162 | I201-I201 | -0.8086 |
| R18-Q456 | -1.5376 | R28-F208 | -0.7707 | E202-I215 | -0.7659 |
| R18-E457 | +4.2487 | R17-E117 | 4.2733 | R205-Y210 | -0.721 |
| R19-E457 | +2.6128 | R17-D118 | -0.9536 | F208-Y210 | -0.7794 |
| I22-L453 | -0.5009 | R17-Q119 | -2.3465 | K216-R403 | -0.838 |
| I22-E457 | -0.5345 | R18-Q119 | -0.8704 | K216-L430 | -0.6005 |
| L25-L449 | -1.3153 | L21-E117 | -0.9154 | K216-T433 | 0.891 |
| L25-A450 | -0.4906 | L21-Q119 | -0.7065 | K216-L434 | -0.9962 |
| L25-L453 | -0.5771 | I227-L252 | -0.9567 | K216-Q437 | 0.6498 |
| L26-K442 | -1.0637 | R230-D246 | -0.5433 | F219-P405 | -1.0655 |
| L26-A450 | -0.6108 | R230-L249 | -1.9307 | F219-L406 | -0.8039 |
| E27-K442 | -0.563 | I231-L249 | -0.9429 | F219-I426 | -0.7923 |
| R28-R403 | -0.5559 | K243-K243 | -1.0073 | F219-L430 | -1.4987 |
| R28-P443 | -0.9804 | P193-Y419 | -0.493 | I220-L434 | -0.8734 |
| D29-R403 | 2.9793 | P193-I422 | -0.8711 | I223-I427 | -1.1139 |
| R29-P433 | -0.4608 | P193-L423 | -0.5436 | T226-L423 | -0.5616 |
| D29-P443 | -0.5332 | P193-I426 | -0.7412 | I227-L423 | -0.7135 |
| K30-Q437 | -0.4506 | L194-L406 | -0.6311 | I227-I427 | -0.5469 |
| T31-Q437 | -0.479 | L194-I409 | -0.7333 | R230-L423 | -0.8867 |
| L33-F438 | -1.4224 | L194-I410 | -1.2977 | K243-D417 | -0.7765 |
| L36-L434 | -0.7343 | L194-L413 | -0.5927 | I410-I410 | -0.7806 |
| L36-Q437 | -0.7667 | L194-Y419 | -0.5994 | L413-L413 | -0.719 |
| L36-F438 | -0.9033 | L194-I422 | -0.6963 | E414-Y419 | 2.5438 |
| R17-R209 | -1.5273 | L194-I426 | -0.929 | E414-I422 | -0.5562 |
| L21-F208 | -0.8245 | I197-L406 | -1.3965 | D417-Y419 | -0.5851 |
| Q24-F208 | -1.6762 | L198-I410 | -0.8805 | Q460-R18 | -0.7292 |
| L25-F208 | -0.9052 | I201-L406 | -0.6622 |  |  |
| R28-E203 | +9.9706 | L198-L198 | -1.0981 |  |  |

**Supplementary Table 2.** Total side-chain VDW interaction energies with the opposing subunit.

| Residue<br>Number | Residue<br>Type | E<sub>VDW</sub><br>(kcal/mol) | Residue<br>Number | Residue<br>Type | E<sub>VDW</sub><br>(kcal/mol) |
| --- | --- | --- | --- | --- | --- |
| 17 | ARG | -0.79 | 227 | ILE | -2.61 |
| 19 | ARG | 2.11 | 230 | ARG | -3.84 |
| 21 | LEU | -3.07 | 231 | ILE | -1.03 |
| 22 | ILE | -2.22 | 234 | HSD | -0.33 |
| 24 | GLN | -1.87 | 243 | LYS | -1.89 |
| 25 | LEU | -4.26 | 246 | ASP | -0.98 |
| 26 | LEU | -2.05 | 249 | LEU | -3.52 |
| 27 | GLU | -0.66 | 252 | LEU | -1.34 |
| 28 | ARG | 4.73 | 403 | ARG | 1.78 |
| 29 | ASP | 0.67 | 405 | PRO | -1.60 |
| 30 | LYS | -1.10 | 406 | LEU | -4.94 |
| 31 | THR | -0.57 | 409 | ILE | -1.15 |
| 33 | LEU | -1.54 | 410 | ILE | -3.04 |
| 36 | LEU | -2.51 | 413 | LEU | -2.07 |
| 113 | GLU | -0.36 | 414 | GLU | 1.65 |
| 117 | GLU | -1.74 | 417 | ASP | -2.10 |
| 119 | GLN | -1.40 | 419 | TYR | -0.53 |
| 191 | ASN | -0.37 | 420 | GLN | -0.35 |
| 193 | PRO | -2.75 | 422 | ILE | -2.40 |
| 194 | LEU | -5.74 | 423 | LEU | -3.33 |
| 197 | ILE | -2.04 | 426 | ILE | -2.80 |
| 198 | LEU | -3.24 | 427 | ILE | -1.87 |
| 201 | ILE | -2.26 | 430 | LEU | -2.08 |
| 202 | GLU | -1.49 | 433 | THR | -0.79 |
| 203 | GLU | 9.90 | 434 | LEU | -3.04 |
| 205 | ARG | -1.33 | 437 | GLN | -1.82 |
| 207 | GLN | -2.45 | 438 | PHE | -2.72 |
| 208 | PHE | -4.99 | 442 | LYS | -2.23 |
| 209 | ARG | -0.90 | 443 | PRO | -2.11 |
| 210 | TYR | -2.62 | 446 | SER | -0.84 |
| 215 | ILE | -1.79 | 449 | LEU | -1.85 |
| 216 | LYS | -1.40 | 450 | ALA | -1.41 |
| 219 | PHE | -5.06 | 453 | LEU | -3.09 |
| 220 | ILE | -0.51 | 454 | ALA | -0.73 |
| 222 | VAL | -0.25 | 456 | GLN | -2.53 |
| 223 | ILE | -1.29 | 457 | GLU | 5.23 |
| 226 | THR | -0.80 | 460 | GLN | -0.78 |

**Supplementary Table 3.** Crystallography statistics.

| Protein, ion | C85A / L194A | C85A / L194A / H234C |
| --- | --- | --- |
| PDB I.D. | 7CVS | 7CVT |
| **Data collection** |  |  |
| Space group | C121 | C121 |
| a, b, c (Å) | 232.9, 96.7, 170.7 | 233.0, 98.3, 172.4 |
| $\alpha, \beta, \gamma$ (°) | 90, 131.7, 90 | 90, 132.0, 90 |
| Resolution (Å)* | 33.96 - 3.005 (3.05 - 3.005) | 34.3 - 2.9 (2.95 - 2.9) |
| $R_{merge}$ | 0.173 (1.842) | 0.126 (2.064) |
| $I / \sigma I$ | 24.1 (1.52) | 36.4 (1.87) |
| Completeness (%) | 96.45 (87.70) | 98.90 (96.00) |
| Redundancy | 5.4 (5.3) | 5.0 (4.7) |
| **Refinement** |  |  |
| Resolution (Å) | 33.96 - 3.005 | 34.3 - 2.9 |
| No. unique reflections | 54754 | 63778 |
| $R_{work} / R_{free}$ | 20.1 / 25.5 | 20.1 / 24.1 |
| No. atoms |  |  |
| Protein / ligand | 13235 / 4 | 13245 / 3 |
| B-factor |  |  |
| Protein / ligand | 111.4 / 204.9 | 115.3 / 192.8 |
| R.m.s. deviations |  |  |
| Bond lengths (Å) | 0.005 | 0.006 |
| Bond angles (°) | 1.06 | 1.18 |
| Ramachandran plot†, | 93.5 / 5.75 / 0.75 | 94.14 / 4.82 / 1.03 |
| Rotamer outliers (%) | 0.00 | 0.79 |
| Clash score‡ | 6.32 | 15.97 |

\*Number in parentheses represent values for the highest resolution shell.
†Percentage of residues in the favored / allowed / disallowed region.
‡Statistics were calculated as described in (Chen et al. 2010).

